# Astroplastic: A start-to-finish process for polyhydroxybutyrate production from solid human waste using genetically engineered bacteria to address the challenges for future manned Mars missions

**DOI:** 10.1101/288746

**Authors:** Xingyu Chen, Syeda Ibrahim, Alina Kunitskaya, Kaitlin Schaaf, Zi Fei Wang, Preetha Gopalakrishan, Maliyat Noor, Harry Wilton-Clark, Jacob Grainger, Alexandra Ivanova, Patricia Lim, Michaela Olsakova, Lalit Bharadwaj, Bilal Sher, David Feehan, Rachelle Varga, Mayi Arcellana-Panlilio

## Abstract

Space exploration has long been a source of inspiration, challenging scientists and engineers to find innovative solutions to various problems. One of the current focuses in space exploration is to send humans to Mars. However, the challenge of transporting materials to Mars and the need for waste management processes are two major obstacles for these long-duration missions.

To address these two challenges a process called Astroplastic was developed that produces polyhydroxybutyrate (PHB) from solid human waste, which can be used to 3D print useful items for astronauts. PHB granules are naturally produced by bacteria such as *Ralstonia eutropha* and *Pseudomonas aeruginosa* for carbon and energy storage. The *phaJ, phaC,* and *phaCBA* genes were cloned from these native PHB-producing bacteria into *Escherichia coli*. These genes code for enzymes that aid in PHB production by converting products of glycolysis and β-oxidation pathways, such as acetyl-CoA and enoyl-CoA, into PHB. To ensure a continuous PHB production system and to eliminate the need for cell lysis to extract PHB, recombinant E. coli was engineered to use the genes in its natural type I secretion system to secrete PHB. The C-terminal of the HlyA secretion tag was fused to phasin (PhaP), a protein originally from *R. eutropha*. Phasin-HlyA electrostatically binds PHB granules and transports them outside of the cell.

In addition to genetically engineering bacteria, a concept for start-to-finish PHB production process was designed. Integrating expert feedback and experimental results, conditions for each step of the process including the collection and storage of waste, volatile fatty acid (VFA) fermentation, VFA extraction, PHB fermentation, and PHB extraction were optimized. The optimized system will provide a sustainable and continuous PHB production system, which will address the problems of transportation costs and waste management for future space missions.

**Financial Disclosure:** Mindfuel Science Alberta Foundation Genome Alberta GenScript Polyferm Canada GeekStarter Alberta Integrated DNA Technologies University of Calgary University of Calgary Cumming School of Medicine University of Calgary Bachelor of Sciences University of Calgary Schulich School of Engineering University of Calgary O’Brien Centre for the Bachelor of Health Sciences City of Calgary Alberta Innovates The funders had no role in study design, data collection and analysis, decision to publish, or preparation of the manuscript.

**Competing Interests:** The authors have declared that no competing interests exist.

**Ethics Statement:** N/A

**Data Availability:** All data are freely available without restriction.

## Introduction

Governments and private enterprises alike are gearing up for travel across our Solar System. Plans to colonize nearby planets are underway, with Elon Musk spearheading the initiative to put a human colony on Mars by 2030. Furthermore, NASA is planning a manned exploratory mission to Mars as soon as the 2030s. Several other space agencies have similar plans and timelines for their own respective Mars explorations. This exciting time in our history nonetheless comes with the challenges of long-term space travel.

Two current ecological and economic challenges for space travel include the sustainable management of waste produced in space and the high cost of shipping materials to space. Recycling of waste on Mars will be paramount because manned missions will need to recover as much water and oxygen as possible to sustain life. Human waste must also be treated to minimize health risks for the crew of a Mars mission. All of this must be accomplished while preserving the natural Martian environment. Transportation of goods is also a major economic barrier for long-duration missions to Mars because the current cost of shipping materials up to low Earth orbit is $22,000 USD per kilogram ($10,000 USD per pound) [1]. This expense will constrain early Mars mission crews in the supplies that they can bring or ship from Earth to Mars and may not allow astronauts to account for every tool they may require during their mission. One way to mitigate this challenge is to develop a system to produce necessary items in space as needs arise from solid organic waste, an abundant resource from space travel.

This project worked on a unique solution to both challenges of future Mars missions; upcycle human waste by using it as a feedstock for *E. coli* engineered to produce bioplastic, which can then be 3D printed into useful tools on-site.

Poly(3-hydroxybutyrate) (PHB), a bioplastic, is produced in nature by many bacterial species. Literature has shown that PHB can be produced using a variety of feedstocks, including glucose and volatile fatty acids (VFAs) [2]. Since human waste contains both glucose and VFAs, it is a potentially useful feedstock for PHB production. The overarching goal for the synthesis component of the project was to produce PHB by utilizing the nutrients present in human waste. To accomplish the efficient conversion of organic feedstocks into PHB, *Escherichia coli* were genetically engineered to produce PHB by manipulating the endogenous beta-oxidation and glycolysis pathway.

The choice of the genes and metabolic pathway was informed by the types of organic compounds available in human fecal waste that can serve as substrates for PHB synthesis. Analysis of solid human waste showed that human fecal waste contains VFAs (short-chain fatty acids such as acetic acid, propionic acid, and butyric acid), long chain fatty acids, and glucose which can be used as substrates for PHB production [3]. Therefore, the synthesis system must make use of a wide range of carbon sources and transform them into the desired product, PHB.

Many pathways and genes were manipulated in this study to use the organic substrates in human fecal waste to synthesize PHB [4, 5, 6, 7]. To utilize the fatty acids found in human fecal waste, the fatty acid beta-oxidation pathway within *E. coli* was manipulated. A construct was designed to convert short-and medium-chain-length VFAs to PHB. This construct contained the *phaJ4* gene from *Pseudomonas putida*, encoding for enoyl-CoA hydratase that converts the enoyl-CoA into (R)-hydroxybutyrate and the *phaC1* gene, a gene from *Pseudomonas aeruginosa* encoding the PHA synthase, which converts the (R)-hydroxyacyl-CoA into polyhydroxybutyrate (PHB) [8]. This production pathway not only uses VFAs but can also use undigested long-chain fatty acids in human fecal waste, thus maximizing the substrates available for PHB synthesis.

Another production system construct contained the *phaCBA* genes, which uses the products of glycolysis for PHB production. The *phaCBA* pathway was adopted from the naturally existing *phaCAB* operon found in *R. eutropha* H16 which is involved in the biosynthesis of PHB [4]. Transcription of the *phaCAB* operon leads to expression of the following enzymes in the order: pha synthase (*phaC*), 3-ketothiolase (phaA), and acetoacetyl-CoA reductase (*phaB*). 3-ketothiolase converts acetyl-coA to acetoacetyl-CoA. These enzymes use acetyl-coA, which results from degradation of carbohydrates and VFAs, as a primary substrate to produce PHB [4]. The acetoacetyl-CoA reductase leads to conversion of acetoacetyl-CoA to (R)-3-hydroxybutyryl-CoA. Finally, pha synthase converts (R)-3-hydroxybutyryl-CoA to PHB [4]. Hiroe et al. showed that the content of PHB is dependent on the expression of *phaB* [4]. Therefore, this construct contained the rearrangement from native *phaCAB* operon to *phaCBA* to promote higher expression of phaB for increased PHB produced.

Codon optimization helps to increase the translational efficiency of gene by modification of DNA sequence of one species into a sequence of codons most recognized another species. As a result, protein expression is maximized in the respective organism. Therefore, the constructs described above were codon optimized to increase their translational efficiency within *E. coli*.

To produce PHB, the synthetic feces (see methods) was prepared as a feedstock for the bacterial cultures. PHB is produced and stored as intracellular granules that range in size from 60-80 nm within the bacteria [9]. This creates a challenging problem: efficient extraction of the desired PHB from inside the cell. After the evaluation of the advantages and disadvantages of different extraction methods in terms of our space application, the Type I hemolysin secretion pathway, which is endogenous to *E. coli*, was selected as a method to extract PHB [9] due to its advantages over traditional extraction methods such as chemical lysis and heat-induced lysis.

Chemical lysis traditionally uses solvent extraction employing chemicals such as chloroform and sodium hypochlorite [10]. Although this method produces efficient yields and is relatively simple, it requires chemicals which need to be transported to and from Mars. This introduces a high cost for the project. In addition, this method requires the manual input of chemicals, which would demand time from astronaut’s schedule. The chemical lysis would kill the cells and eliminate the continuous production of PHB. In addition, the cellular debris will make the separation of the PHB more difficult. Heat-induced lysis was another method for PHB extraction which depends on the T4 lysis genes (holins and endolysins) under the control of the pR Lambda promoter with thermosensitive repressor proteins cI587. When heated, lysis genes become activated [11]. This system is advantageous because it does not require chemicals for lysis. However, the energy consumption from the heating of fermenters is an issue because energy is another limited resource in space. In addition,heat-induced lysis also kills the bacterial cells, not compatible with the goals of continuous and sustainable production and extraction system. Given the disadvantages associated with these conventional PHB extraction methods, the BBa_K2018024 from the SDU-Denmark 2016 was as the basis for the design of a Type I hemolysin secretion mechanism.

PHB is secreted through the hemolysin pathway that is endogenous to *E. coli*. Naturally, the hemolysin toxin (HlyA) is secreted from *E. coli* as a defence mechanism. This is a single-step process that involves three additional proteins: HlyB, HlyD, and TolC. HlyB is an active transport protein that uses ATP in the cytoplasmic membrane, HlyD is a membrane fusion protein that spans the periplasm and connects the inner and outer membrane proteins, and TolC rests in the outer membrane. When HlyB recognizes the C-terminus of HlyA, it stimulates formation of the secretion channel and the toxin is secreted [12].

To use this system in our biologically engineered cells, the C-terminus of HlyA, which is endogenous to *E. coli*, was added to the end of a phasin molecule. Phasin, coded for by *phaP*, is a small structural protein found in bacteria that naturally produce PHB. It electrostatically binds to intracellular PHB, and therefore the PHB granules with bound phasin-HlyA tag are secreted as one by type 1 secretion machinery. Granules secreted through this mechanism range from 20-60 nm in size [9].

Part BBa_K2018024 from the SDU-Denmark 2016 team was used for the design of a Biobrick for PHB secretion. This part contains a coding region for *phasin (phaP*), originally from *R. eutropha*, as well as a coding region for the HlyA secretion tag (*hlyA*), originally from *E. coli*. The entire sequence for *E. coli* was sequence-optimized to improve protein expression in the chassis, *E. coli* BL21(DE3), and restriction sites were removed to make it compatible with all iGEM RFC assembly standards. An IPTG-inducible T7 bacteriophage promoter was chosen to upregulate the production of Phasin-HlyA because increased production of Phasin has been shown to decrease PHB granule size, which could increase the efficiency of PHB translocation across the cellular membrane [13]. In addition, genes under the control of an inducible promoter have much higher rate of transcription than those that are constitutive. The *E. coli* BL21(DE3) genome contains a coding region for T7 RNA polymerase under the control of an IPTG-inducible promoter, and it was therefore chosen as an ideal chassis. Furthermore, a FLAG tag (amino acid sequence DYKDDDDK) was added to the N-terminus of the Phasin-HlyA fusion protein for easier isolation of our protein during protein expression analyses.

Process design: In the first step of the process, astronaut’s feces are collected into a storage tank using a vacuum toilet. Feces are then transferred into another tank and left to ferment for three days with natural gut flora to increase the concentration of volatile fatty acids (VFAs) that are later consumed by engineered *E. coli* to produce PHB. Next, the liquid containing VFAs is separated from solid particles using centrifugation and filtration. The resulting liquid containing VFAs is then passed to another storage tank. From there, VFAs are added to a fermenter inoculated with the engineered PHB-producing and PHB-secreting *E. coli*. Lastly, the resulting PHB is extracted from the liquid harvest stream. The PHB can then be used in a Selective Laser Sintering (SLS) 3D printer without the need for additional processing [14].

During the development of the project, the team consulted experts to determine the most appropriate project application. When the feasibility of this project was evaluated, there was four possibilities of where to integrate PHB production: wastewater treatment plants, small-scale wastewater treatment plants in developing countries, landfill leachate treatments, and on Mars colonies with human solid waste as feedstock. The former three project applications were deemed unfeasible due to high up-front costs. After careful consideration, the team chose Mars colonies as the final project application.

From conversations with experts, such as Dr. Robert Thirsk and Col. Chris Hadfield (two Canadian astronauts), the team was alerted to considerations for designing systems for space. The team thus considered In-Flight Maintenance Requirements, where any machine that is brought to and used in space must be one that can be easily repaired by crew members. This aspect in design was taken into consideration by Process Development in designing the key components of the PHB-producing system. The team also made sure that the system was compliant with current policies and regulations, such as the treaties released by the United Nations Committee on the Peaceful Uses of Outer Space (COPUOS).

Based on COPUOS recommendations, the team emphasized containment of foreign biological systems in their project design. Dr. Nicole Buckley (a microbiologist at the Canadian Space Agency (CSA)) provided the team with possible safety concerns with the PHB-producing system, such as viral shedding from feces that can increase disease prevalence on board the spacecraft. To address this issue, the team is currently considering the implementation of UV treatment to assist in sterilizing the human waste after it has been fermented with natural gut flora. Dr. Buckley also indicated the possibility of having abnormal or decreased growth rates of bacteria in microgravity. To address this, it was decided that the PHB-producing system would be implemented under the gravitational pull of Mars in a colony. Furthermore, Dr. Buckley also expressed the difficulty of bring cell maintenance materials to space, which pushed the team towards the less material-intensive PHB secretion system, as opposed to a lysis system.

Dr. Matthew Bamsey (the Chief Systems Engineer at the German Aerospace Center) was also consulted; he provided NASA documents on Equivalent System Mass, which provides an universal method of calculating the required energy and efficiency of items brought to space, and mentioned that the team should consider the NASA Life Support Baseline Values and Assumptions, which is a document containing a set of values to be used in the Equivalent System Mass Analysis. Current applications of solid waste in space were also accounted for on the recommendation of Dr. Pascal Lee (the Principal Investigator of the Haughton-Mars Project at the NASA Ames Research Center).

Made in Space, Inc. is the company that developed and implemented the installation of a 3D printer currently in use on the ISS. Dr. Derek Thomas is the Senior Materials Scientist at this company and offered the team valuable advice for safety and encouraged the team to explore the flammability, off-gassing, and other physical characteristics of PHB on Mars. Secondary processing may also be a method to obtain desired properties not found in the unrefined PHB. Moreover, although Dr. Thomas did not foresee any major concerns with the use of powder PHB in a SLS 3D printer, it may be difficult to control the powder in microgravity. This is another reason why the team chose to apply the system on a Mars colony, where gravitational pull is present. The feasibility of using PHB as 3D printing feedstock was also supported by Dr. Bruce Ramsay, the founder and CTO of Polyferm, a commercial PHB production company.

## Materials and Methods

### Genetic Constructs

#### Codon Optimization

Genes of interest were codon optimized for use in *E. coli*. This was done by breaking the DNA sequence of nucleotides into codons, which were replaced with codons of higher frequency in E.coli that translates to the same amino acids. The genes were optimized using the codon usage table for *E. coli* [15].

#### Competent *E. coli* BL21 (DE3) and DH5 Preparation

The *E. coli* BL21 (DE3) and *E. coli* DH5α cells used in the experiments of this project were obtained from Dr. Sui-Lam Wong and Dr. Richard Moore of University of Calgary, respectively. The protocol used to transform both strains of *E. Coli* was provided by Dr. Richard Moore.

The strain to be transformed was cultured in 2 mL LB broth and shaken at 28℃ overnight. The strained was then subcultured 1:50 into 50 mL LB with 10 mM MgSO4 and 1 mM KCl and shaken at 28℃ until OD600 reaches 0.3 to 0.4. After chilling on ice for at least 10 minutes, the mixture was transferred to a 50 mL pre-chilled tube. The mixture was then centrifuged at 2,500g for 8 minutes at 4℃ and was resuspended in 10 mL ice-cold 100 mM CaCl2. The cells were gently mixed on ice and incubated on ice, for a period of time between 10 minutes to overnight. Then, the tube of cells was centrifuged at 2500xg for 8 minutes at 4℃ and resuspended in 500 μL 100 mM CaCl2 and 10% glycerol. The mixture was mixed by gently pipetting up and down and left on ice for at least 10 minutes. The cells were then separated to 1.5 mL pre-chilled tubes as 50 μL aliquots, while remaining on ice.

The competency of the cells was then assessed by transformation with pUC19 from NEB. 50 μL of competent cells and 1μL pUC19 (50 pg/μL) were combined and left on ice for 40 minutes. The mixture was then put into a water bath at 37℃ for 45s. On ice, 400 μL of LB were added and shaken for 1 hour at 37℃. 225 μL of the culture was then streaked on an ampicillin (100µg/mL) plate and grown overnight at 37℃.

### Cloning

The IDT geneBlocks were digested and ligated to pET29b(+). The restriction enzymes used are given in Table 1. After the genes were inserted in the plasmid, *E. coli* DH5α was transformed with the vector. DH5α strain was used because of its ability to produce a high copy number of plasmids. Plasmid miniprep was carried out and the obtained plasmids were digested and run on gel electrophoresis to confirm if ligation was successful.

**Table 1.**
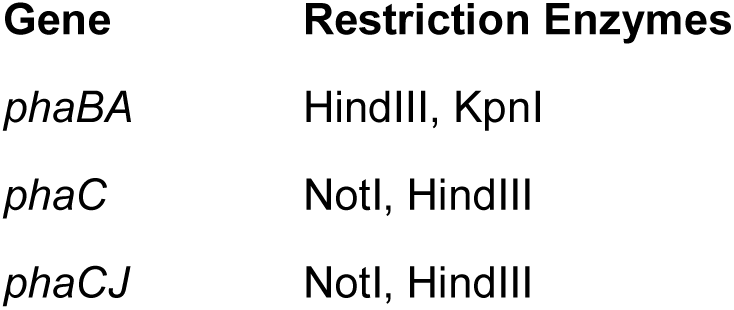
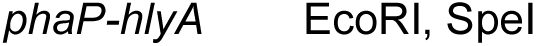
Restriction enzymes used for digestion of pET29b(+) and the respective genes.

After cloning the genes given in Table 1, some of the genes were ligated together. Digested *phaBA* and *phaC* vectors were excised from low-melting point gel and ligated together followed by transformation of *E. coli* DH5α with the resulting plasmid. The restriction enzymes used were XbaI and HindIII. Standard protocols for digestion, ligation, and transformation were followed. After confirmation whether the genes were successfully ligated, the final operons (*phaCBA* and *phaCJ*) were sequenced.

After cloning the genetic constructs in *E. coli* DH5α and confirmation of the gene sequences, plasmid miniprep was carried out. The obtained plasmids were then used to transform *E. coli* BL21 (DE3) because of this strain’s ability to synthesize more protein. Hence, all genetic constructs were cloned in *E. coli* BL21 (DE3), which was used for experiments.

### Digestion of constructs and vectors

In order to carry out a restriction digest, DNA of known concentration was obtained. In a 1.5 mL microcentrifuge tube, 1μg DNA suspended in TE buffer or ddH2O was added. For a 20 μL solution, 1 μL of the desired restriction enzyme(s) was added to the tube. 2 μL of 10x buffer for the respective enzyme was added. Finally, ddH2O was added to the tube to adjust final volume to 20 μL. The tube was then allowed to incubate at 3°C for an hour followed by heat inactivation of the restriction enzymes by placing the tube in water bath at 80°C for 20 minutes. The tube was then stored in −20°C.

The restriction enzymes used for our experiments were from New England Biolabs (NEB) and the manufacturer’s protocol was used for the respective restriction enzymes.

### Ligation of Vector and DNA insert

In order to clone the respective DNA used for experiments, the DNA parts were ligated to a plasmid backbone (pSB1C3 or pET29b(+)). The vector was then used to transform *E. coli* DH5ɑ for plasmid propagation or *E.coli* BL21(DE3) for protein expression. The protocol used for ligation experiments were obtained from NEB. In a 1.5 mL microcentrifuge tube, 50 ng of the digested vector DNA suspended in TE buffer or ddH2O was added. For a 20 μL reaction, 2 μL of 10x T4 DNA ligase buffer was added to the tube. The DNA insert:vector ratio used was 3:1 and the desired amount of plasmid was calculated and the appropriate volume of vector was added to tube. 1 μL of T4 DNA ligase was added to tube followed by incubation of tube on ice placed at room temperature overnight. Finally, the tube was stores at −20°C for use in further experiments.

### Plasmid Miniprep

Plasmid miniprep was carried out to analyze the DNA. Cells were cultured in culture tubes containing LB and the respective antibiotic. 2 mL of the overnight culture was transferred by pipetting into a 2 mL microcentrifuge tube. The microcentrifuge tube was spun at 3500xg for 10 minutes and a cell pellet was obtained. The supernatant was discarded. Cells were then resuspended in 300 μL resuspension buffer. 300 μL of lysis buffer was added and the tube was inverted gently. The tube was left to incubate at 23℃ for 3-5 minutes. 300 μL of precipitation buffer was added and tube was inverted gently. The resulting white suspension was centrifuged at 14,000xg for 10 minutes at 23℃. The supernatant obtained was transferred to a clean 1.5 mL microcentrifuge tube using pipette. 650 μL of isopropanol was added and tube was gently inverted. It was then allowed to incubate at 23℃ for 10 minutes. The tube was placed in microcentrifuge and spun at 14,000xg for 10 minutes at 4°C. The resulting supernatant was discarded and pellet was washed with 500 μL cold 70% ethanol. The tube was placed in micro Centrifuge at 14,000xg for 5 minutes at 4°C and supernatant was discarded. The pellet was dried by placing microcentrifuge in speed vac for 15-30 minutes, or until no more liquid was visible in the tube. The pellet was suspended in ddH₂O and stored at −20°C.

### Bacterial Transformation

In order to test our genetic constructs, *E. coli* BL21(DE3) was transformed with the respective genes inserted in pET29b(+) or pSB1C3. 50 μL aliquot of competent *E.coli* BL21(DE3) cells were obtained from −80 C freezer and placed on ice to thaw just before use. 0.3-1 μg of DNA was added to cells using pipette and the microcentrifuge tube containing the cells was gently flicked to allow DNA to mix with cells. CaCl₂ concentration was adjusted to maintain competency of the cells and the mixture was incubated on ice for 45 minutes. The cells were then heat shocked for 60-75 seconds at 42°C. The microcentrifuge tube was placed on ice for 5 minutes. 2 mL of SOC medium was added to the aliquot of cells followed by incubation for 60-90 minutes at 37°C, shaking at 200 rpm. 50-100 μL of re-suspended culture was obtained and plated on agar plate with appropriate antibiotic. The cells were spread using aseptic technique. Finally, the agar plate was placed in incubator set to 37°C overnight or until desired growth was observed.

### Confirmation Digest

After transforming *E. coli* with the desired vector containing gene of interest, a confirmation digest was carried out to analyze the presence of DNA insert in the vector. The four conditions used for the confirmation digest were: (i) Plasmid of interest containing vector digested with restriction enzymes used for ligation, (ii) Plasmid with no insert digested with restriction enzymes used for ligation, (iii) Plasmid of interest containing vector digested with restriction enzyme that cuts at the region of DNA insert and another enzyme that cuts the backbone, and (iv) Plasmid with no insert digested with restriction enzyme that cuts at the region of DNA insert and another enzyme that cuts the backbone. Finally, the product of the reaction was analyzed using agarose gel electrophoresis. A 1% agarose gel was run at 100V for 30 minutes.

### PHB Synthesis

#### Utilizing Acetyl-CoA to produce PHB

Synthetic feces (see below for Preparation of Synthetic Feces and Supernatant) were fermented to produce acetyl-CoA for PHB synthesis. After fermentation of synthetic feces, glucose is converted to acetic acid followed by acetyl-CoA formation. To test whether *phaCBA* construct can synthesize PHB using fermented synthetic feces, three replicates of overnight cultures having OD600=0.4-0.8 of each negative control (vector with no insert) and vector with *phaCBA* insert were made. The overnight cultures were grown in LB media and the respective antibiotic (kanamycin, 50µg/mL). In 125 mL Erlenmeyer flasks, 10 mL overnight cultures were added followed by the addition of 10 mL fermented synthetic feces, 100 uL MgSO4 (2mM), 5 uL CaCl2 (0.1 mM), 10 ml M9 salts (1x), and 5 uL 1 M IPTG. Final volume in flasks were adjusted to 50 mL with ddH2O. The flasks were covered with aluminum foil and placed in a shaker for 24 hours at 37°C and 100 rpm. Solutions in the flasks were obtained next day and poured into 50 mL Falcon™ tubes. Chemical lysis was carried out using sodium hypochlorite extraction to analyze PHB production.

#### Utilizing VFAs to produce PHB

Other VFAs present in fermented synthetic feces are propionic acid, butyric acid, etc. Beta-oxidation pathway uses the VFAs and *phaCJ* construct can be used to convert the products to produce PHB in bacteria. Thus, four different conditions were used to test whether PHB is synthesized using fermented synthetic feces: negative control (vector with no insert), positive control (vector with insert and glucose present in media), pure VFAs, and fermented synthetic feces. Pure VFAs were used to test how much PHB is produced by utilizing VFAs when no glucose is present. 9 replicates of overnight cultures of *E. coli* BL21(DE3) transformed with pET29b(+) containing phaCJ insert were made along with 3 replicates of overnight cultures of the negative control (pET29b(+) with no insert). OD600 of the overnight cultures was adjusted to lie between 0.4 and 0.8. 10 mL of the cultures were added to 125 mL Erlenmeyer flasks. 100 uL MgSO4 (2mM), 5 uL CaCl2 (0.1 mM), 10 ml M9 salts (1x), and 5 uL 1M IPTG were added to each flask.

7 mL of 20% glucose solution was added to the positive control. Propionic acid, acetic acid, and butyric acid were in 3:2:1 ratio to the pure VFAs condition. Lastly, 10 mL of fermented synthetic feces was added to each of the remaining replicates. The flasks were covered with aluminum foil and placed in a shaker for 24 hours at 37°C and 100 rpm. The cultures were obtained the next day and poured into 50 mL Falcon™ tubes. The tubes were centrifuged, and the resulting supernatant was discarded. Cell pellet was resuspended in 1x PBS for extraction and the respective OD600 were recorded before proceeding to sodium hypochlorite extraction.

#### Sodium Hypochlorite Extraction of PHB

Sodium hypochlorite (bleach) was used to chemically lyse the bacterial cells, causing them to release PHB into their media, which can then be isolated via centrifugation. This extraction method of carried out for simplistic analysis of PHB production. 50 mL of PHB-producing *E. coli* BL21(DE3) overnight cultures were centrifuged at 3275xg for 10 minutes in a 50 mL Falcon™ Tube. The supernatant was discarded, and the pellet was resuspended in 5 mL 1X PBS solution. The resuspended mixture was centrifuged again at 3275xg for 10 minutes. The supernatant was discarded, and the pellet was resuspended in 5 mL 1% (v/v) Triton X-100 in PBS and left to incubate for 30 minutes at 23°C. The mixtures were centrifuged at 3275xg for 10 minutes. The supernatant was discarded. The pellet was resuspended in 5 mL 1X PBS solution and centrifuged at 3275xg for 10 minutes. The supernatant was discarded.

The mixture was then resuspended in 5 mL of sodium hypochlorite and left to incubate at 30°C for 1 hour. After the hour, the mixture was centrifuged at 3275xg for 20 minutes. The supernatant was discarded, and the pellet was washed in 5 mL 70% ethanol. This wash step was repeated several times, with 20 minutes of centrifugation at 3275xg between each step. The extracted PHB powder was then allowed to dry overnight in an open tube.

#### PHB Secretion

*E. coli* BL21(DE3) transformed with *phaCAB-phasin-HlyA* in a pSB1C3 plasmid backbone was used to test the production and secretion of PHB into the extracellular media. *E. coli* BL21(DE3) transformed with *phaCAB* in a pSB1C3 plasmid backbone was used as a negative control. All experiments were carried out in triplicates. 1 mL of overnight culture was inoculated in 49 mL LB media + 3% glucose + 30 µg/mL chloramphenicol (total volume of 50mL), induced with 5 µL 1 M IPTG, then incubated at 37°C and 150 rpm in an aerobic environment for 48 hours. At 24-hour intervals, the culture was transferred to a 50mL Falcon™ tube and 500 µL of 1M CaCl2 (0.05549g CaCl2) was added to to the cultures to reach a final concentration of 0.01 M CaCl2. After 10 minutes incubation at 23℃, the solution was centrifuged at 50xg for 5 minutes and the supernatant removed to a new 50mL Falcon™ tube. This pellet contains the secreted fraction of PHB and cellular debris. The supernatant was centrifuged at 3270xg for 10 minutes and the supernatant discarded. This pellet contains the intracellular PHB granules inside live cells, which were not secreted.

To remove cellular debris, the pellet of secreted PHB was resuspended in 5 mL of 1% Triton X-100 in PBS, incubated for 30 minutes at 23℃, then Centrifuged at 3275g for 10 minutes. The supernatant was discarded and the pellet was dried overnight in an open tube at 23℃. To extract PHB from the intracellular fraction, the pellet was treated with sodium hypochlorite and dried overnight in an open tube at 23℃. This protocol was adapted from Rahman et al. 2013.

## Characterization of PhaJC and PhaCAB

### SDS-PAGE

Overnights cultures of *E. coli* containing the pet29b(+), pet29b(+)-phaCJ, and pet29b(+)-phaCBA vector were made and left to incubate at 37℃ in a shaker for 16 hrs at 200 rpm. 100μL of the overnight cultures were then subcultured in a new tube of 2mL LB for 2hr. The bacteria were then induced with 0.1mM IPTG and left in the shaker for another 4 hours. The cultures were then pelleted in microcentrifuge tubes spun at max. The pellets were put in the −20℃ freezer and left overnight. The next day, the pellet was thawed at room temperature. The pellet was resuspended in 100μL of STET buffer. 1mg of lysozyme was added to the resuspension and left to incubate in a water bath at 37℃ for 15 minutes. The suspensions were then sonicated for 5 seconds on and 5 seconds off while on ice. Centrifugation was used to separate the soluble and insoluble proteins by spinning the sonicated suspensions at max for 10 min. The soluble portion was pipetted into another microcentrifuge tube. The insoluble pellet was resuspended in 50μL loading buffer. Protein loading dye was made by combining 50μL of loading buffer with 3μL DTT. 15uL of the soluble and insoluble protein fractions were aliquoted into microcentrifuge tubes. 3uL of the loading dye was added to each protein sample. The protein samples with the dye were boiled for 5 minutes to denature the proteins and then ran on a 1X SDS-PAGE gel for 30 mAmp for 30-40 minutes.

The materials used for the gel and procedures for setting up an SDS-PAGE gel are listed below.

1x SDS gel loading buffer

- 50mM tris-Cl (pH 6.8)
- 100mM dithiothreitol
- 2% sodium dodecyl sulfate
- 0.1% bromophenol blue
- 10% glycerol

1x Tris-Glycine electrophoresis buffer:

- 25mM tris
- 250mM glycine
- 0.1% (w/v) sodium dodecyl sulfate

Stacking gel:

- dH_2_O
- 30% acrylamide mix
- 1.0M tris (pH 6.8)
- 10% sodium dodecyl sulfate
- 10% ammonium persulfate
- TEMED

10% Resolving gel:

- dH_2_O
- 30% acrylamide mix
- 1.5M tris (pH 8.8)
- 10% sodium dodecyl sulfate
- 10% ammonium persulfate
- TEMED

HPLC PHB quantification was conducted using high performance liquid chromatography (HPLC) by digesting samples in concentrated acid to produce crotonic acid. The protocol was based on the methodology outlined in [16]. PHB-containing samples were first digested in 1 ml concentrated sulphuric acid in a 90 degrees Celsius water bath for 30 minutes. After the digestion was complete the samples were cooled on ice and 4 ml of 0.014N sulphuric acid was rapidly mixed in. Samples were diluted using 0.014N sulphuric acid, with dilution factors ranging from 10-100. The samples were filtered through a 0.2 micron filter before running them on a Aminex HPX-87H ion-exclusion HPLC column to measure crotonic acid. PHB content in the samples was calculated by assuming an 60% conversion, which was found by digesting known concentrations of standard industrial-grade PHB samples.

## PHB Extraction

### Chemical coagulation of PHB

One of the design options we considered for PHB extraction used the principle of chemical coagulation. In order to test chemical coagulation of PHB particles we first simulated nanoscale PHB, which was the expected size range of secreted PHB [9]. 5 g/L industrial-grade PHB in suspension in distilled water was sonicated in 250 ml batches in a probe-type sonicator to break up the particles. The suspension was then allowed to settle for 48 hours before the top phase was separated by decanting. Three conditions were tested with replicates to show that adding a coagulant helped the nanoparticles to agglomerate into larger particles that could be more easily recovered by centrifugation. The test conditions included, centrifugation at 1000 RPM, centrifugation at 3750 RPM, and centrifugation at 3750 RPM after making up a 10mM calcium chloride concentration in the sample. Amount of PHB extracted was estimated by measuring the absorbance of each of the sample at 600nm.

### Electrocoagulation of PHB

We also tested electrocoagulation to extract PHB nanoparticles from a suspension in distilled water and from synthetic feces supernatant. Supernatant from the synthetic feces recipe was prepared by centrifuging samples in 50 ml tubes at 3750 RPM for 20 minutes, removing the supernatant and then centrifuging the supernatant again at 3750 RPM for 20 minutes. 5 g/L suspension of PHB in distilled water was prepared by sonication the suspension in 250 ml batches in a probe-type sonicator. The suspension was then allowed to settle for 48 hours before the top phase was separated by decanting. The electrocoagulation cell was prepared by connecting an iron anode and a steel cathode to a 9 volt battery and placing the test sample in the cell [17]. Three types of samples were tested: PHB suspension in distilled water to demonstrate whether PHB particles could be made to settle using electrocoagulation, synthetic feces supernatant as a negative control, and a 1:1 mixture of synthetic feces supernatant and PHB suspension to test whether electrocoagulation could work to separate PHB from feces supernatant. The last test sample simulated the real mixture PHB would have to be separated from in the process designed for Mars. Each cell was run for 3 hours after 250 ml of the test sample was loaded into the cell. Qualitative results were obtained by observing the settlement.

## Preparation of Synthetic Feces and Supernatant

Due to safety and ethical concerns with using human feces, synthetic feces mimicking the chemical and physical properties of real feces were used for the experiments. In addition to water, human feces contain fats, carbohydrates, nitrogenous material, minerals, and bacterial debris. Synthetic feces were prepared using a modified recipe from National Aeronautics and Space Administration (NASA). The original recipe included water, dry baker’s yeast, microcrystalline cellulose, psyllium, miso paste, oleic acid, sodium chloride, potassium chloride and calcium chloride (Table 2).

**Table 2.** Original NASA recipe for preparation of synthetic feces [18].

| Compound | Amount (g per kg feces) |
| --- | --- |
| Water | 800 |
| Baker's yeast (dry) | 60 |
| Microcrystalline cellulose | 30 |
| Psyllium | 35 |
| Miso paste | 35 |
| Oleic acid | 40 |
| NaCl | 4 |
| KCl | 4 |
| CaCl2 | 2 |

The recipe was modified to include yeast extract in place of dry baker’s yeast to avoid fermentation of synthetic feces by yeast and oleic acid was removed from the recipe due to challenges with processing samples containing oils on the HPLC (Table 3).

**Table 3.** Modified recipe used for preparation of synthetic feces

| Compound | Amount (g per kg feces) |
| --- | --- |
| Water | 800 |
| Yeast extract | 60 |
| Microcrystalline cellulose | 35 |
| Psyllium | 35 |
| Miso paste | 35 |
| NaCl | 4 |
| KCl | 4 |
| CaCl<sub>2</sub> | 2 |

Some experiments required synthetic feces supernatant, where solid particles were removed. To obtain synthetic feces supernatant, synthetic feces were centrifuged at 3000 rcf for 20 min. The liquid portion was then collected and sterilized using 0.20 μm syringe filters. The solid portion was discarded. The supernatant was stored at −20°C, if required.

## Effect of Temperature on VFA Fermentation

Bench scale experiments were performed to determine the optimal operating temperature for the VFA fermentation step of the process. The two temperatures selected for the experiment were 37°C and 22°C corresponding to the optimal bacterial fermentation temperature and expected room temperature in a Mars habitat, respectively. Synthetic feces were fermented with *E. coli* BL21(DE3) transformed with pET29b(+) vector without any inserts (control), PHB-producing *E. coli* transformed with a PHB-producing part from iGEM Imperial College 2013 (“Imperial condition”) and PHB-producing *E. coli* transformed with PHB-producing part from iGEM Tokyo 2012 (“Tokyo condition”) inside a shaker at 80 rpm and desired temperature. Different fermentation durations were tested including 3 and 5 days. Synthetic feces were inoculated with 1 mL of overnight *E. coli* cultures per 100 mL of synthetic feces. Erlenmeyer flasks used for fermentation were sterilized before the experiment and were covered with aluminum foil during fermentation. After fermentation, fermented synthetic feces were centrifuged at 3000 rcf for 20 minutes. The supernatant was then collected. A sample of the supernatant was taken for VFA measurements using titrations, while the remaining supernatant was sterilized using 0.20 μm syringe filters. Sterilized supernatant was then fermented with PHB-producing bacteria, as previously described. The amounts of PHB produced from each VFA fermentation condition were then compared.

## VFA Quantification using Titrations

A titration method described by Anderson and Yang (1992) was used to quantify VFA concentration in synthetic poop supernatant. To quantify VFA, 50 mL of diluted sample were placed in a 100 mL beaker [19]. The beaker was placed on the magnetic stir plate and a magnetic stir bar was used to continuously mix the sample during titrations. A pH probe was placed into the solution to measure the change in pH. After recording the initial pH, 1N sulfuric acid was added in 100 μL −1 mL increments until a set point of 5.1 followed by 3.5. The volume of acid titrated to reach the desired pH set point was recorded. The following equations were used to calculate the concentration of VFA:

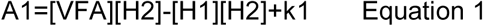

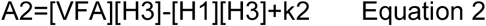

Where A1= molar equivalent of acid consumed to the first titration point

> A2= molar equivalent of acid consumed to the second titration point
>
> [H1],[H2], [H3] = concentration of H+ ions initially and at the two titration points
>
> [VFA]= concentration of VFA
>
> k1= conditional dissociation constant for carbonic acid
>
> k2= combined dissociation constant for VFA

Equations above can be rearranged to solve for VFA concentration.

## Equivalent System Mass Analysis

Equivalent System Mass (ESM) analysis, a method recommended by NASA to evaluate advanced life support systems, was used to evaluate the feasibility of the proposed process and to compare different process options [20’]. ESM analysis is used to identify which of the proposed designs has the lowest launch cost based on mass, volume, power, cooling, and crewtime needs in place of traditional dollar costs. The following formula was used to determine ESM value for the proposed process:

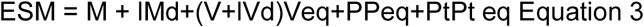

Where ESM = equivalent system mass (kg)

> M = mass of the system (kg)
>
> IM = logistics mass (kg/day)
>
> d = mission duration (days)
>
> V = volume of the system (m3)
>
> IV = logistics volume (m3/ day)
>
> Veq= volume equivalency factor (kg/m3)
>
> P = power requirement of the system (kW)
>
> Peq= power equivalency factor (kg/kW)
>
> Pt = thermal control power requirement (kW)
>
> Pt eq= thermal control equivalency factor (kg/kW)

The crew time requirements were not considered in the analysis due to lack of data. ESM equivalency factors were obtained from NASA’s Baseline Values and Assumptions for Advanced Life Support Systems [21] and are summarized in Table 4.

**Table 4.** Equivalency factors for volume and power used in ESM analysis of the process

| ESM Equivalency | Factor Units | Nominal |
| --- | --- | --- |
| Shielded volume | kg/m | 3216.5 |
| Unshielded volume | kg/m | 39.16 |
| Power | kg/kW | 87 |
| Thermal control | kg/kW | 146 |

## Process Development Assumptions

A number of assumptions were made based on Anderson et al. (2015) when developing the physical process that starts with feces and end with a bioplastic powder that can be used in a 3D printer [21]. The nominal surface habitat duration was assumed as 600 days with a crew size of 6 people. The process was designed to accommodate a maximum of 150 g of fecal matter per crew member per day corresponding to a volume of 150 mL per crew member per day. Based on the amount of fecal matter, the predicted water recovery for VFA and PHB production was assumed to be 123 g or 123 mL per crew member per day. The power required for the process will be supplied using solar panels and nuclear sources, which were assumed to be available on Mars.

## Results

### Successful cloning of *phaCJ* and *phaCBA*

A double digest confirmation was carried out to confirm the presence of *phaCJ* insert in pET29b vector. The digestion utilized NotI and HindIII restriction enzymes. The length of *phaJ* is 474 bp. Hence, the total sum of *phaC* and *phaJ* genes is around 2.2 kb. The products of confirmation digest were analyzed through agarose gel electrophoresis. The results showed the presence of two linear DNA of sizes 5.4 kb and 2.2 kb. The different lanes show results of confirmation digest on plasmid miniprep of cell cultures prepared from different colonies, which were obtained after transformation of *E. coli.*

The pet29b(+) vectors containing *phaCBA* insert were analyzed through gel electrophoresis after performing digestion confirmation. Fig 2. shows the results of digesting pET29b vector containing *phaCBA* operon using KpnI and NotI. The length of *phaC, phaB*, and *phaA* genes are 1.8 kb, 759 bp, and 1.2 kb. The products of the confirmation digest were analyzed through agarose gel electrophoresis. The results showed the presence of DNAs of length 5.3 kb, 2.8 kb, and 1.0 kb respectively. The length of pET29b vector is around 5.4 kb and the sum of the length of *phaCBA* operon is 3.8 kb. The sum of the two lower bands was observed to be 3.8 kb.

**Fig 2.**
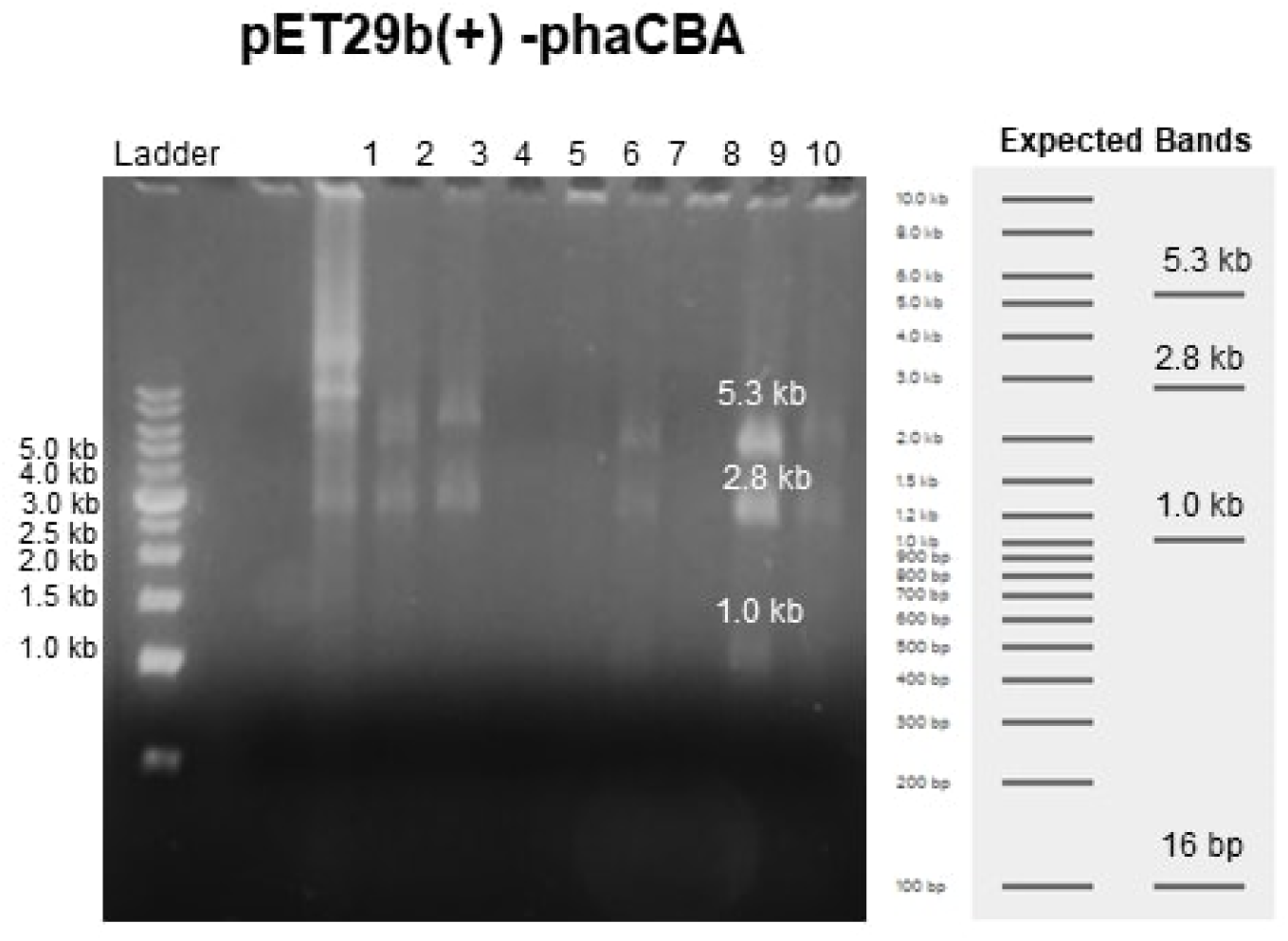
*phaCBA* was successfully cloned into the pet29b(+) vector *E. coli*. Double digest confirmation of pet29b(+)-phaCBA with NotI and KpnI ran on 1% agarose gel.

An SDS-PAGE gel was ran to determine whether the *E. coli* cloned with the vectors containing *phaCJ* and *phaCBA* could produce the proteins of interest. 0.1mM IPTG was added to a 2mL subculture of overnight culture cultivated for 16h and further cultivated for 2h. The soluble and insoluble protein fractions were loaded in the lanes outlined in Fig 3. *phaJ* and *phaC* activity was detected in the insoluble protein fraction. A dark band was observed in the pET29b(+)-phaCJ lane between 17-22 kDa, which is in reasonable agreement with the approximate mass of the predicted PhaJ product (17.7kDa). Another protein was observed in the insoluble portion of pET29b(+)-phaCBA, which is in reasonable agreement with the mass of the predicted PhaA product (41.4kDa). These protein bands were not seen in the control pET29b(+) lanes. The protein bands of the PhaC (65.1kDa) and PhaB (27.2) for pET29b(+)-phaCBA were not observed. The regions where the expected protein bands for PhaC from pet29b(+)-phaCJ and PhaC and PhaB from pet29b(+)-phaCBA were smeared, unclear, and had many close bands together.

**Fig 3.**
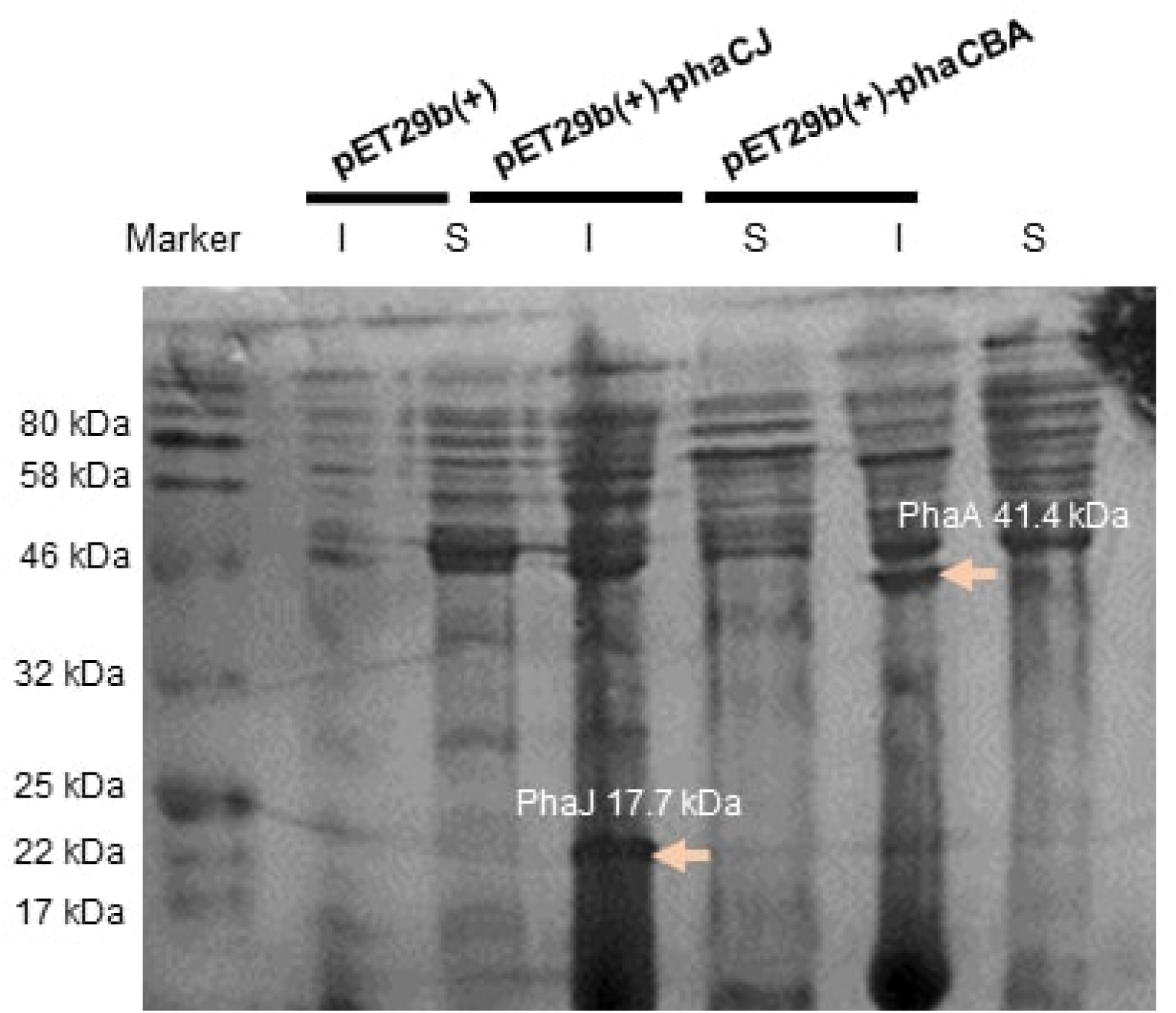
SDS-PAGE of *E. coli* containing vectors pet29b(+)-phaCJ and pet-29b(+)-phaCBA show bands of PhaJ and PhaA. SDS-PAGE of soluble (S) and insoluble (I) proteins obtained from *E. coli* containing pET29b(+)-phaC1J4 induced with 0.1mM of IPTF at 37°C. Cells were lysed with lysozyme, sonicated, and centrifuged to divide the proteins into a soluble and insoluble fraction. The proteins were denatured after boiling. The gel was ran at 30mA for 50 minutes and stained with Coomassie Blue.

To confirm the ability of *E. coli* transformed with our pET29B(+)-phaC1J4 construct to produce PHB, overnight cultures were grown for 24 hours and induced with 0.1 mM of IPTG while supplementing them with chemical media mimicking fermented synthetic poop supernatant (FSPS). Cells were then left to culture for 16 hours. To collect the resulting PHB, sodium hypochlorite extraction was performed.

**Fig 4.**
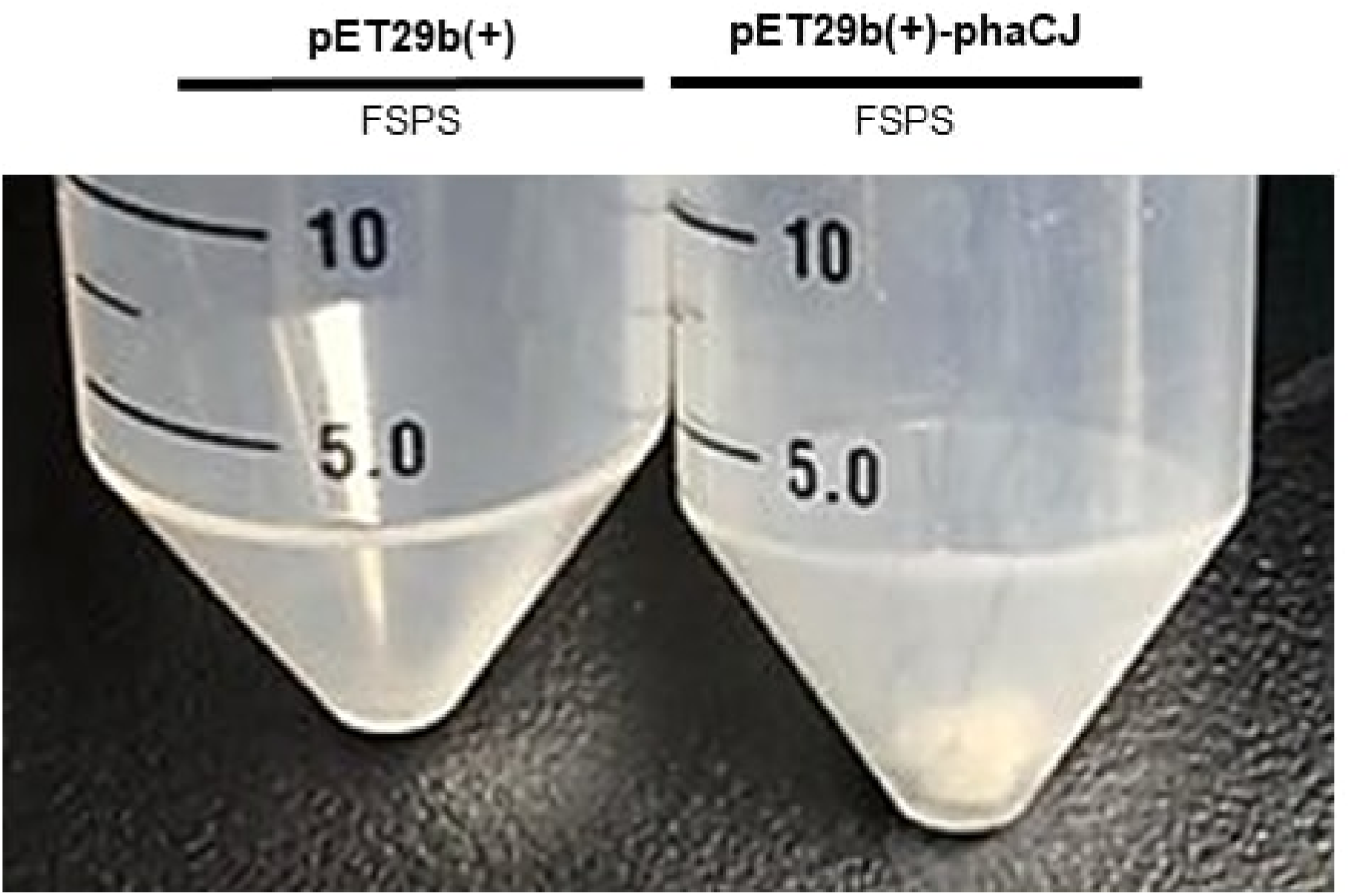
PHB extracted from *E. coli* containing pet29b(+)-phaCBA using the sodium hypochlorite extraction method. PHB extracted from phaC1J4-expressing cells cultured for 24 hours and supplemented with fermented synthetic poop supernatant (FSPS) for 16 hours. PHB was extracted using TritonX-100, sodium hypochlorite, and ethanol in a series of washes and incubation. The negative control tube contained *E. coli* transformed with the pET29B(+) vector.

To confirm the ability of *E. coli* transformed with our pET29B(+)-phaC1J4 construct to produce PHB, overnight cultures were grown for 24 hours and induced with 0.1 mM of IPTG while supplementing them with chemical media mimicking fermented synthetic poop supernatant (FSPS). Cells were then left to culture for 16 hours. To collect the resulting PHB, we performed sodium hypochlorite extraction.

**Fig 5.**
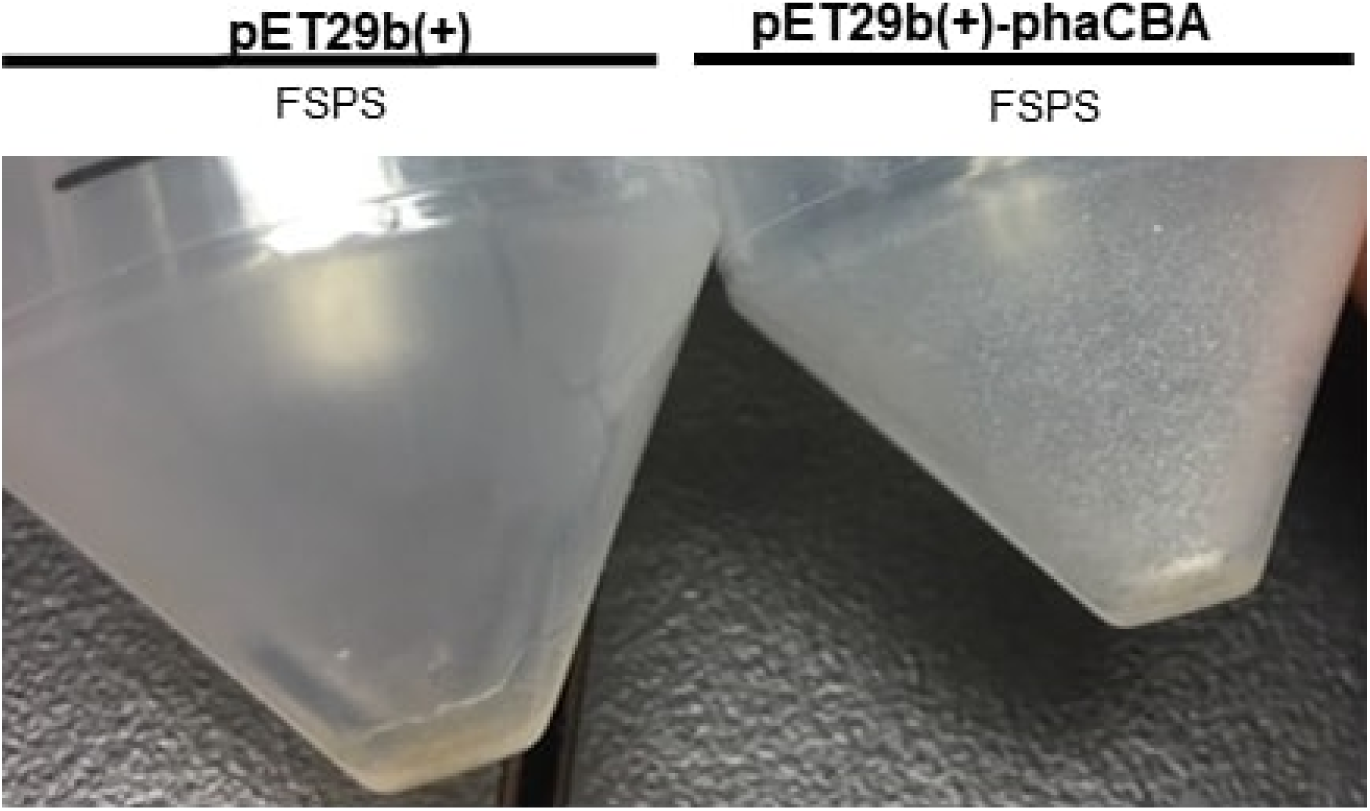
PHB extracted from *E. coli* containing pet29b(+)-phaCBA using the sodium hypochlorite extraction method. PHB extracted from phaCBA-expressing cells cultured for 24 hours and inoculated with FSPS for 16 hours. PHB was extracted using TritonX-100, sodium hypochlorite, and ethanol in a series of washes and incubation. The negative control tube contained *E. coli* transformed with the pET29b(+) vector.

## PHB Characterization Using High-performance Liquid Chromatography (HPLC)

### phaCBA

The product extracted from *E. coli* BL21(DE3) transformed with vector containing the *phaCBA* operon was digested with sulfuric acid using the method in section 1. The sample was then analyzed using HPLC to detect if crotonic acid was present. The HPLC results showed that crotonic acid peak was observed. A standard curve was also generated by using known concentrations of industrially-produced PHB to determine the conversion of PHB to crotonic acid (results not shown). Hence, the amount of crotonic acid was used to determine the amount of PHB in sample. By using the predicted conversion of 80% from PHB to crotonic acid [16], the amount of crotonic acid detected was found to be 0.0282006 mM using a dilution factor of 32.

### phaCJ

Similarly, the product extracted from *E. coli* BL21 transformed with vector containing the *phaCJ* operon was digested with sulfuric acid. The sample was then analyzed using HPLC to detect if crotonic acid was present. There were two different digestion times used: 20 mins and 30 mins and low and high dilution factor for each digestion time. All samples showed a peak for crotonic acid indicating that PHB was present. However, the amount of crotonic acid detected was too low to estimate the percentage of PHB in sample.

**Fig 7.**
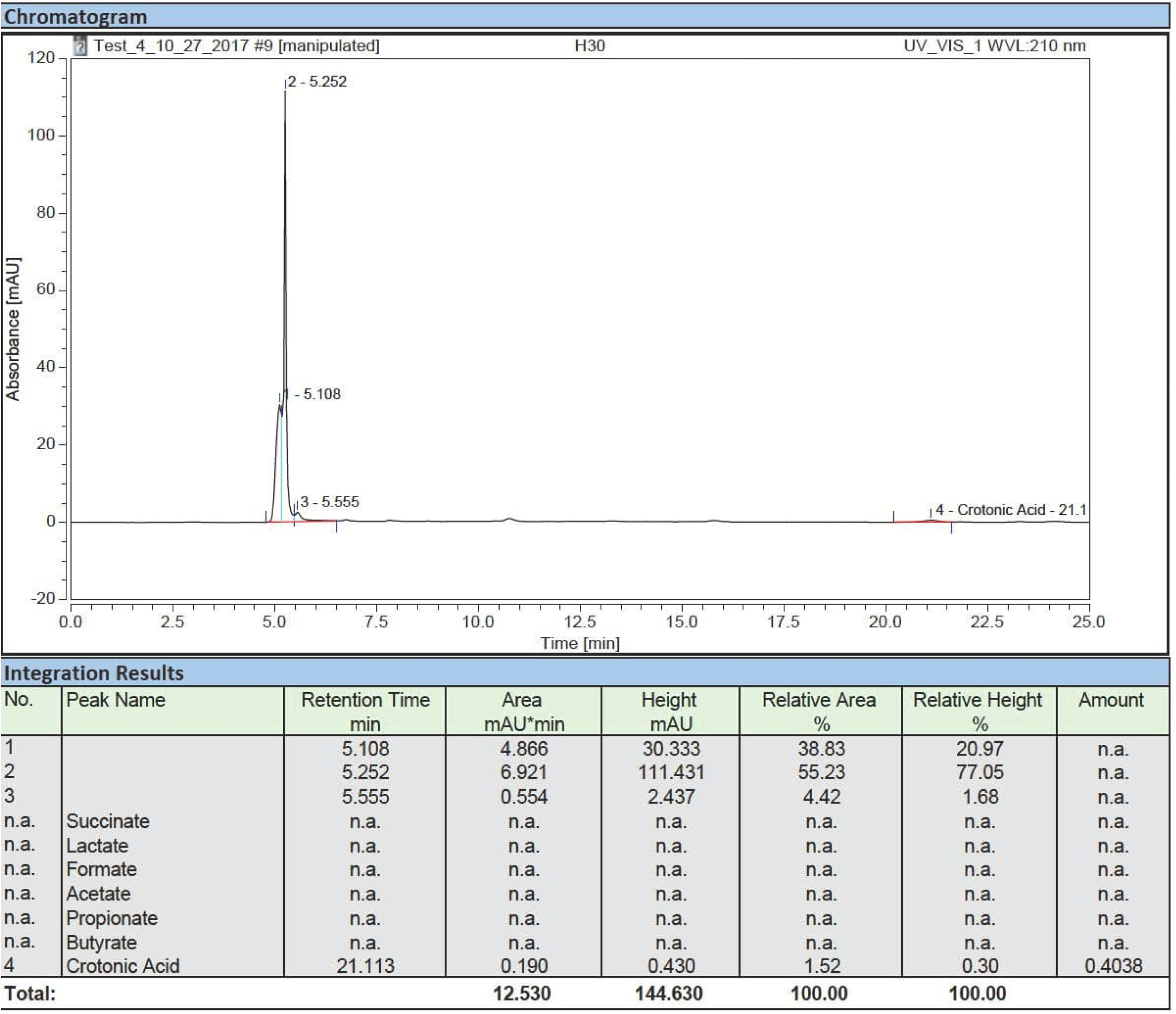
HPLC results of sample containing PHB digested with sulfuric acid HPLC analysis of PHB produced by *E. coli* containing pet29b(+)-phaCJ. The PHB sample was obtained from sulfuric acid digestion of product extracted from *E. coli* (BL21) transformed with *phaCJ* construct. Product was digested for 30 mins and a dilution factor of 30 was used.

## Successful cloning of phasin-HlyA and phaCBA-phasin-HlyA

The successful cloning of the pSB1C3 vector with phasin-HlyA into *E. coli* DH5ɑ was demonstrated by digest confirmation with EcoRI-HF and SpeI. The length of the phasin-HlyA gene is 889 bp and the length of the pSB1C3 vector is 2.0 kb. DNA bands of these sizes were obtained when the products of the confirmation digest were analyzed through agarose gel electrophoresis (Figure 8).

**Fig 8.**
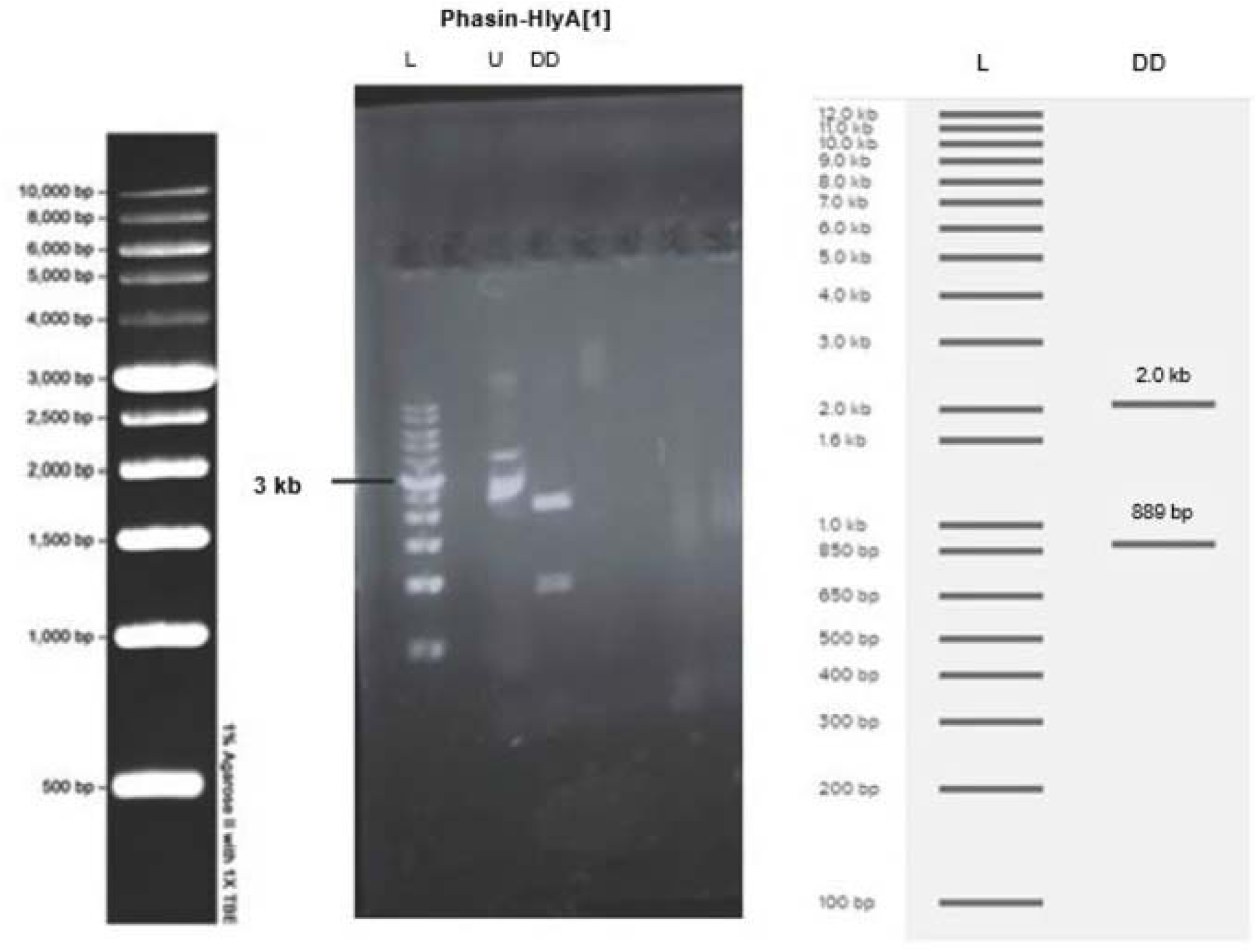
Screening results of DH5α transformed with phasin-HlyA in a pSB1C3 vector. (DD) Plasmid from the colony was digested with EcoRI-HF and SpeI then run on a 1% agarose gel at 100V for 30 minutes. DNA bands around 889 bp and 2.0 kb are visible in lanes 3-4. (L) The molecular ladder is visible on the far left and the expected band sizes, obtained from Benchling Virtual Digest, are visible on the right. (U) Undigested plasmid was used as a control.

To create a PHB-secreting strain of *E. coli*, *phaCAB* was ligated into a pSB1C3 vector already containing phasin-HlyA. The successful cloning of the pSB1C3 vector with phaCAB-phasin-HlyA into *E. coli* DH5ɑ was demonstrated by digest confirmation with NotI-HF and XbaI. The length of the phaCAB and phasin-HlyA genes ligated together is 4.8 kb and the length of the pSB1C3 vector is 2.0 kb. DNA bands of these sizes were obtained when the products of the confirmation digest were analyzed through agarose gel electrophoresis (Fig 9). Two colonies ([5] and [6]) were successfully cloned, however colony [6] was used for all secretion assays.

**Fig 9.**
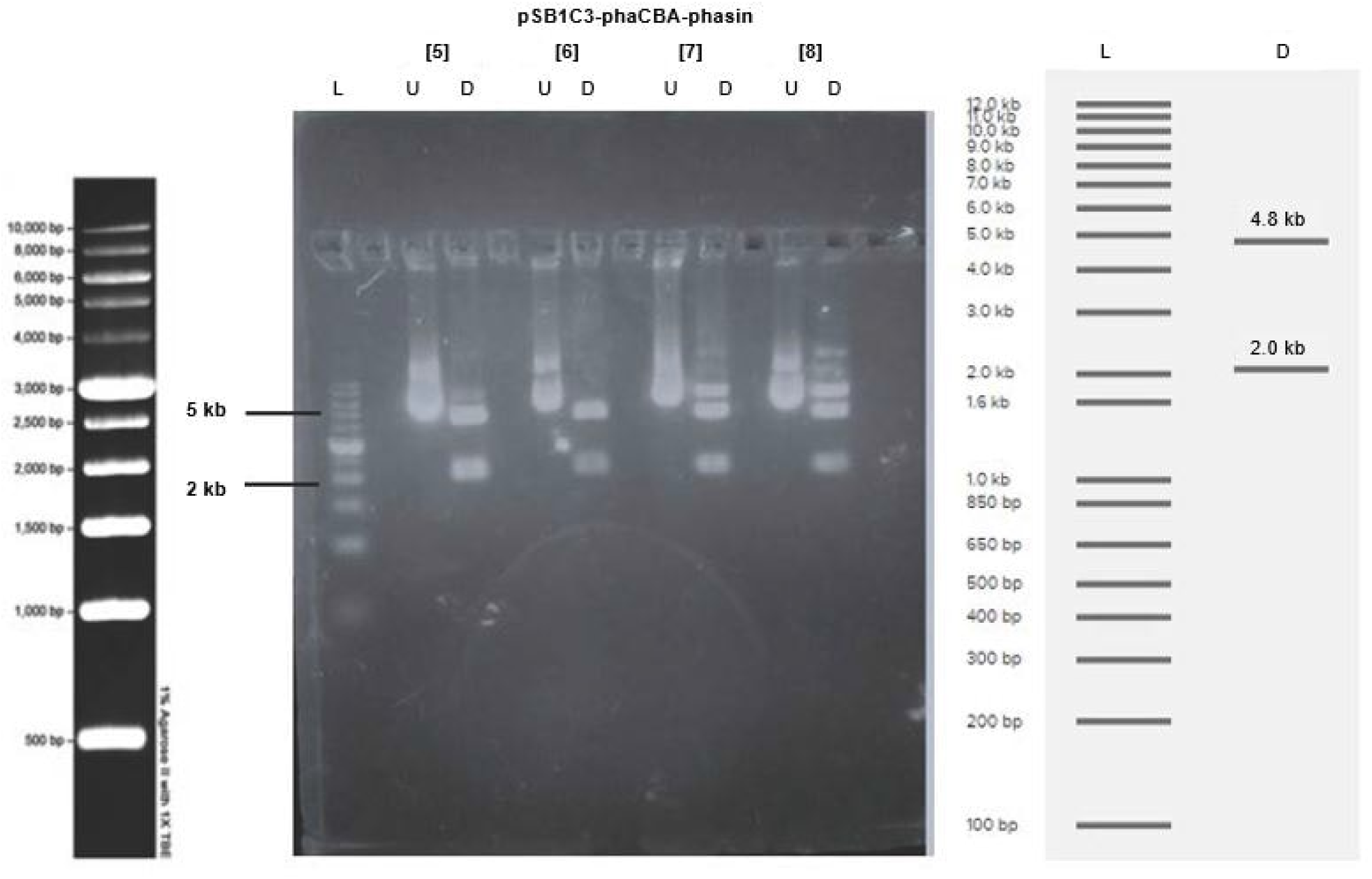
Screening results of DH5 α transformed with pha-CAB-phasin-HlyA in a pSB1C3 vector. (D) Plasmid from the cells were digested with NotI-HF and XbaI then run on a 1% agarose gel at 100V for 30 minutes. DNA bands around 4.8 kb and 2.0 kb are visible in lanes 4 and 7, from colony [5] and [6] respectively. (L) The molecular ladder is visible on the far left and the expected band sizes, obtained from Benchling Virtual Digest, are visible on the right. (U) Undigested plasmid was used as a control.

## Secretion Assay

The mass of PHB present in secreted and extracellular fractions of *E.coli* BL21(DE3) after the secretion assay was obtained by measuring the mass of each 50mL Falcon™ tube before each trial and after the final washing and drying of the PHB step. The mass of the secreted PHB pellet was corrected by subtraction of 0.05549g to account for the CaCl2 that was added during extraction. After 48 hours of incubation at 37℃ there was a 114% increase in the amount of PHB secreted by cells that contain pSB1C3-phaCAB-Phasin-HlyA compared to the negative control (cells with pSB1C3-phaCAB only). However, after 24 hours of incubation at 37℃ there was very little difference in the amount of PHB secreted by the experimental and control groups (Fig 10).

**Fig 10.**
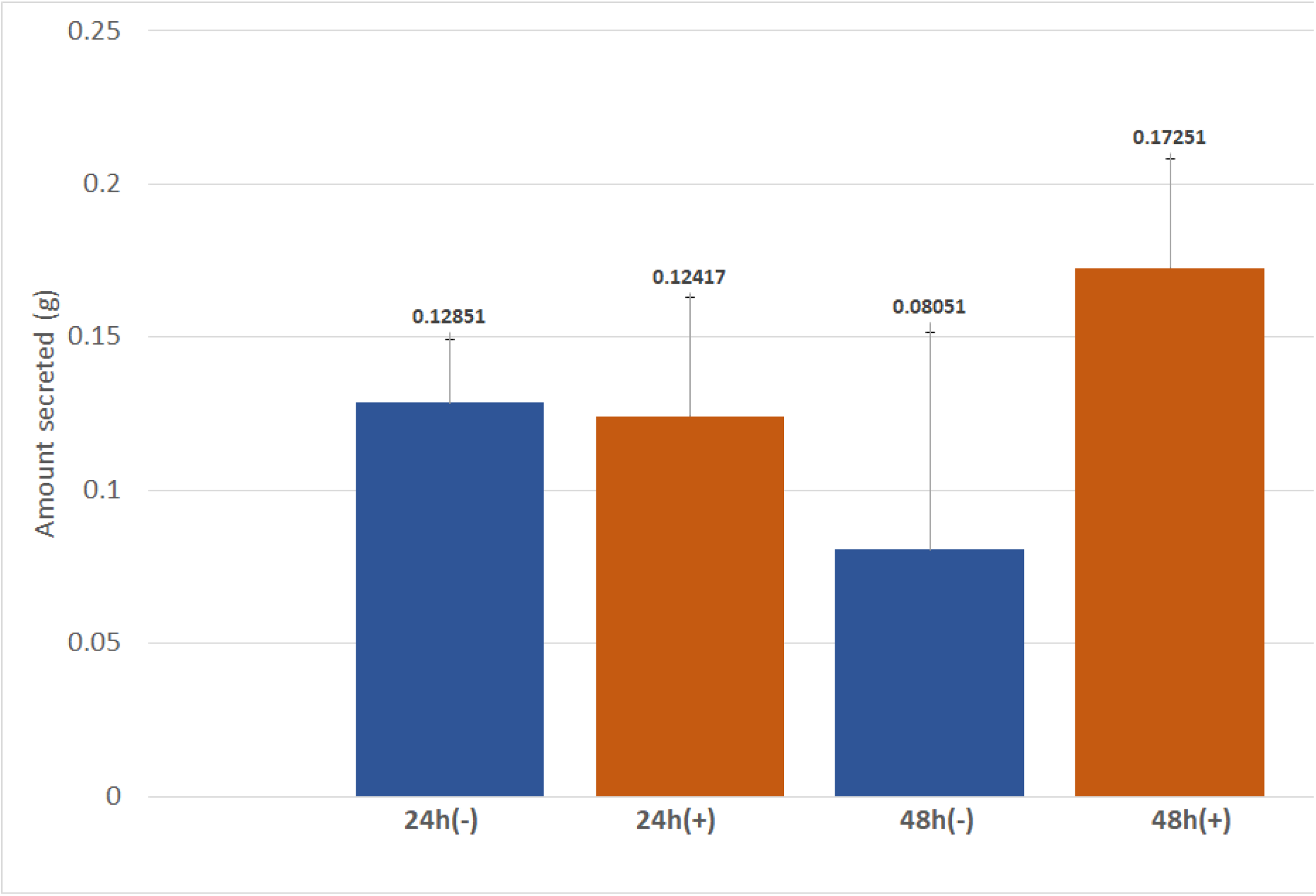
Secretion Assay Results. Amount of PHB secreted (g) after *E. coli* BL21(DE3) transformed with pSB1C3-PhaCAB-Phasin-HlyA (pSB1C3-Phasin) or pSB1C3-PhaCAB (-) were induced then incubated in 50mL of LB (30 µg/mL chloramphenicol) + 3% glucose for 24 or 48 hours. Each condition was carried out in triplicates.

## Process development

### Effect of Temperature on VFA Fermentation

The first VFA fermentation experiment showed higher VFA production at 37°C than at 22°C after 3 days of fermentation (Fig 11). The reported results are an average of 3 titrations performed for each sample.

**Fig 11.**
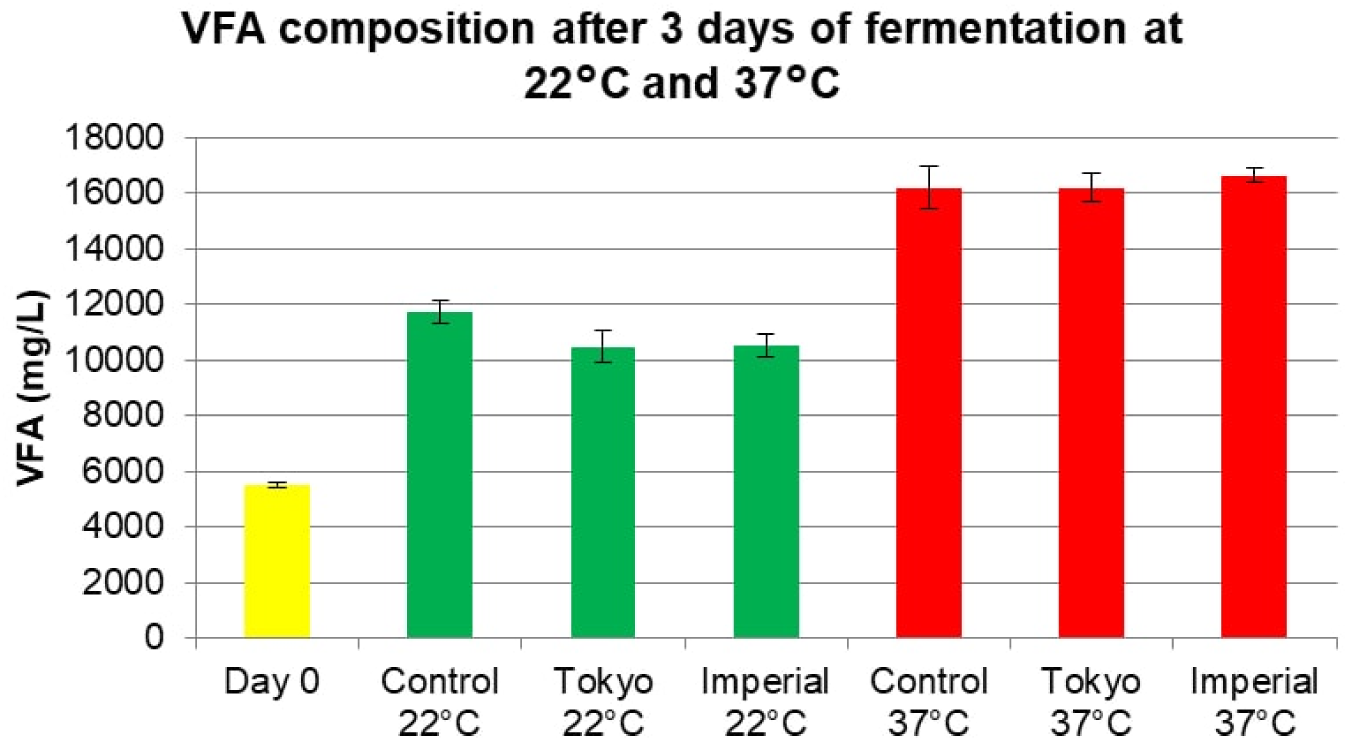
VFA concentrations after 3 days of fermentation at 22°C and 37°C. Amount of VFA (mg/L) in the synthetic feces supernatant after a 3 day fermentation at 80 rpm and desired temperatures with *E. coli* BL21(DE3) transformed with pET29b(+) vector without any inserts (control), PHB-producing *E. coli* transformed with a PHB-producing part from iGEM Imperial College 2013 (“Imperial condition”) and PHB-producing *E. coli* transformed with PHB-producing part from iGEM Tokyo 2012 (“Tokyo condition”). Error bars represent standard deviation for 3 measurements.

The second VFA fermentation experiment confirmed higher VFA production at 37°C than at 22°C (Fig 12). In addition to a 3-day fermentation (denoted as D3), a 5-day fermentation (denoted as D5) was also introduced for this experiment. Higher VFA concentrations were observed on day 5 of fermentation. The reported results are an average of 3 titrations performed for each sample.

**Fig 12.**
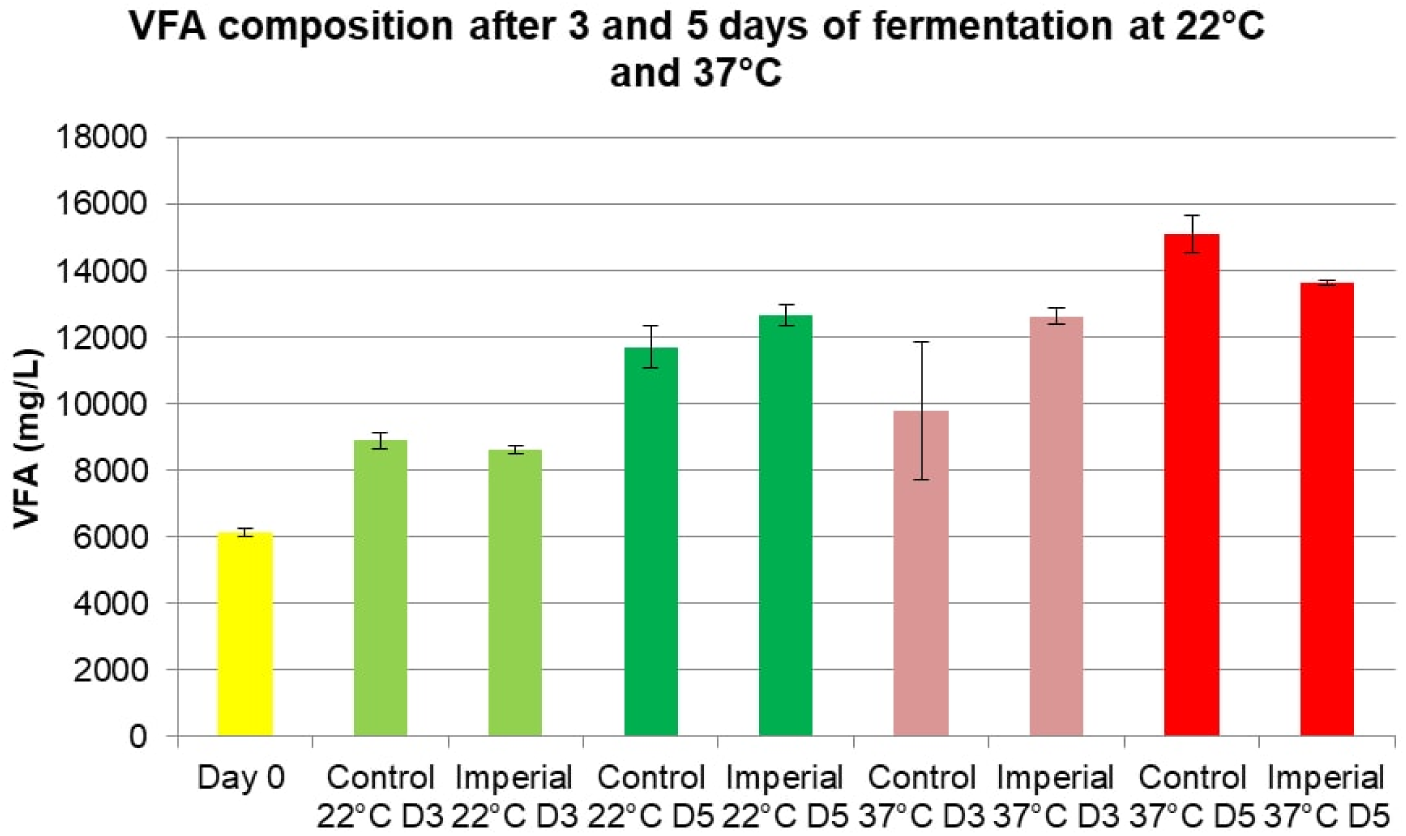
VFA concentrations after 3 and 5 days of fermentation at 22°C and 37°C. Amount of VFA (mg/L) in the synthetic feces supernatant after 3 and 5 day fermentation at 80 rpm and desired temperatures with *E. coli* BL21(DE3) transformed with pET29b(+) vector without any inserts (control), PHB-producing *E. coli* transformed with a PHB-producing part from iGEM Imperial College 2013 (“Imperial condition”). Error bars represent standard deviation for 3 measurements.

Next, fermented synthetic feces supernatant was sterilized and cultured with PHB-producing bacteria for 2 −3 days at 37°C. PHB-producing bacteria cultured in the supernatant from synthetic feces fermented at 37°C resulted in little to no PHB produced due to little to no bacterial growth, which is believed to be a consequence of low pH due to high concentration of VFA. Although 22°C resulted in lower VFA concentrations, it was selected as the preferred temperature for VFA fermentation due to optimal pH for bacterial growth.

## Equivalent System Mass Analysis

The proposed process was divided into four main stages: collection and fermentation of feces to increase the concentration of VFA (“VFA Fermentation” step), extraction of VFA (“VFA Extraction” step), fermentation to produce PHB (“PHB Fermentation” step), and extraction of PHB from the harvest stream (“PHB Extraction”). The total calculated ESM value accounted for each step of the process as well as intermediate storage tanks and pumps (Table 5).

**Table 5.**
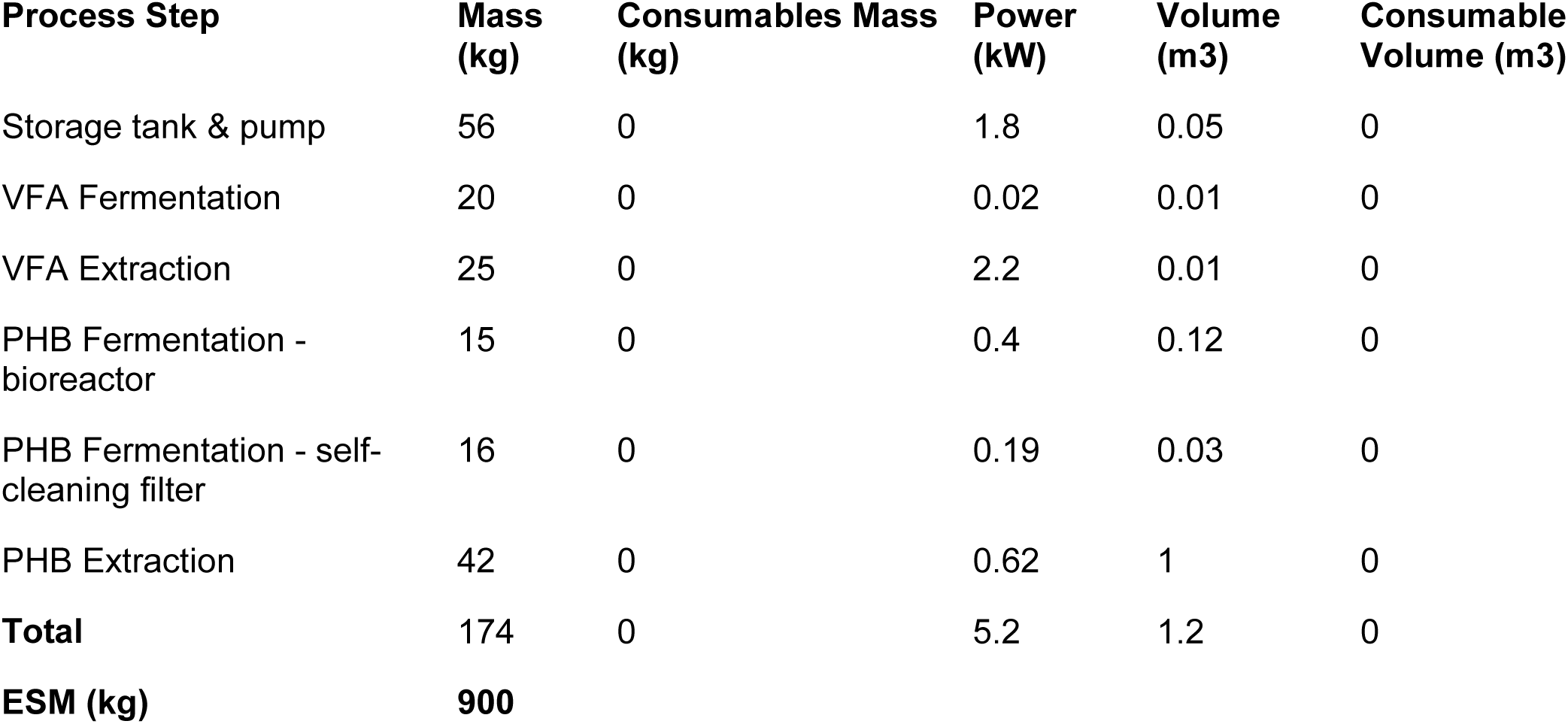
Total ESM value calculated for the proposed process.

| Process Step | Mass (kg) | Consumables Mass (kg) | Power (kW) | Volume (m3) | Consumable Volume (m3) |
| --- | --- | --- | --- | --- | --- |
| Storage tank & pump | 56 | 0 | 1.8 | 0.05 | 0 |
| VFA Fermentation | 20 | 0 | 0.02 | 0.01 | 0 |
| VFA Extraction | 25 | 0 | 2.2 | 0.01 | 0 |
| PHB Fermentation - bioreactor | 15 | 0 | 0.4 | 0.12 | 0 |
| PHB Fermentation - self-cleaning filter | 16 | 0 | 0.19 | 0.03 | 0 |
| PHB Extraction | 42 | 0 | 0.62 | 1 | 0 |
| **Total** | **174** | **0** | **5.2** | **1.2** | **0** |
| **ESM (kg)** | **900** |  |  |  |  |

The total ESM value for our process is lower than some of the systems currently used on the International Space Station. For example, the Water Processor (WP) unit on the International Space Station has an ESM value of about 5,000 kg [22]. After comparing our process to existing life support systems on the International Space Station, we concluded that our process is feasible.

## Chemical coagulation of PHB

We demonstrated in the lab that adding calcium chloride and then centrifuging increased the amount of PHB removed from a suspension in distilled water. We carried out centrifugation at various speeds and then collected absorbance data for the samples to estimate relative quantity of PHB remaining in suspension. The lower the absorbance reading, the higher the amount of PHB that was removed. Absorbance readings and the standard deviations of the various conditions tested are shown in the Fig 13.

**Fig 13.**
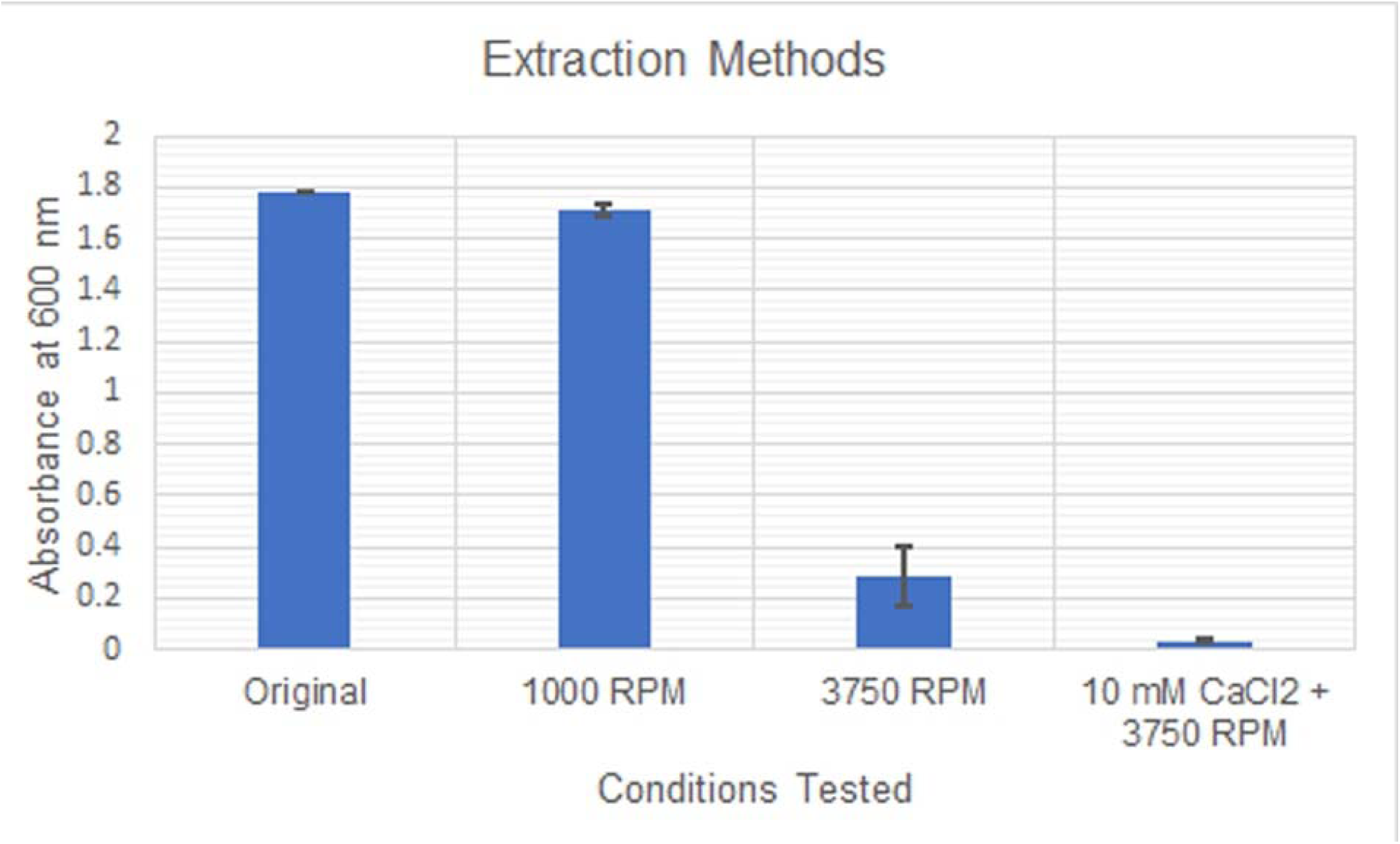
Absorbance readings and their standard deviations for the various extraction methods tested.

As expected, centrifugation at higher RPM and addition of calcium chloride resulted in greater removal of PHB from suspension.

## Electrocoagulation of PHB

From our preliminary experiments with just PHB particles in suspension in distilled water, we were able to demonstrate that PHB particles could settle out via electrocoagulation. A layer of PHB was observed at the bottom of the electrocoagulation cell after it was run for 3 hours. However, a few hours after the cell was shut down, a thin layer of brown powder settled on top of the white layer of PHB. We hypothesized that this was probably iron(III) hydroxide, formed from the excess iron ions released by the anode that did not bind to the PHB.

From our experiments with synthetic feces supernatant, we found that a layer of brown sludge settled at the bottom each time. Even with a 1:1 mixture of PHB suspension and synthetic feces supernatant there was no discernable layer of PHB within the sludge. A sample of the sludge was washed with dilute acid to remove the metal salts that might have been present. However, we were still unable to separate PHB from the sludge.

The electrocoagulation experiments led us to conclude that while it was possible to settle out PHB using electrocoagulation, this method was not suitable for our media which contained a number of salts that interfered with the coagulation process and caused the formation of sludge.

## Discussion

### Synthesis

#### Construct

*phaCJ* and *phaCBA* were successfully cloned into pET29b(+) vectors as shown in the results (Fig 1., Fig 2.) The gel electrophoresis bands of the digested vectors were compared with the expected bands generated from Benchling. Fig1. showed 2 distinct bands for each lane that contained plasmids extracted from various colonies of *E. coli* transformed with the pET29b(+)-phaCJ plasmid. The band sizes were approximately 5.4kb and 2.2kb, corresponding to the predicted bands. Fig 2. verified the successful cloning of pET29b(+)-phaCBA. The expected bands were seen at 5.3kb, 2.8 kb, and 1.0kb. The expected bands image showed an additional band 16bp in size, but this band was not seen in our experimental electrophoresis gel. This is likely because 16bp is a minute size for the DNA examined. Therefore, the DNA of this size may have run off the gel before the current was stopped. However, another gel would have to be run at a shorter time to detect the presence of this 16bp DNA band. Overall, the gel electrophoresis provided sufficient evidence of the successful ligation of the two genes.

**Fig 1.**
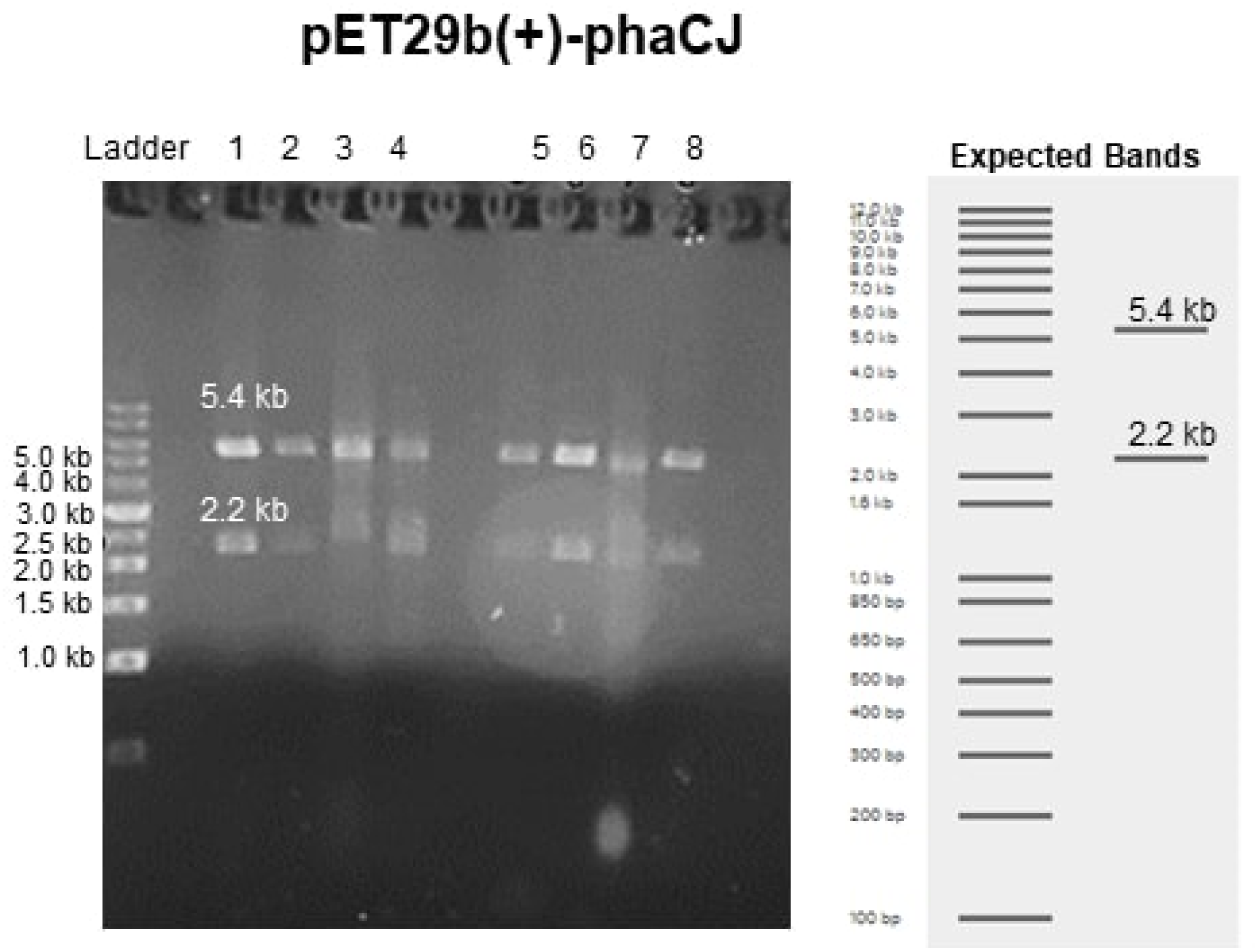
pet29b(+)-phaCJ plasmid digested with NotI and HindIII ran on 1% agarose gel.

#### Enzyme Production

The SDS-PAGE results in Fig 3. showed clear bands for PhaJ (17.7 kDa) and PhaA (41.4 kDa), indicating the expression of the genes from the plasmids. These protein bands were in the insoluble protein fraction. This finding contrast past literature that has found PhaJ in the soluble protein fraction for SDS-PAGE [23]. However, McCool and Cannon found that polyhydroxyalkanoates tend to localize to inclusion bodies, providing an explanation to why these proteins are seen in the insoluble protein fraction [24].

*phaJ* was part of the pet29b(+)-phaCJ plasmid and the *phaA* was part of the pet29b(+)-phaCBA plasmid. The identification of other bands for the corresponding proteins in the same expression plasmid was difficult to identify due to lack of clarity and the closeness of the bands in the regions of interest. Although it is reasonable to assume that the other proteins were also produced alongside the proteins detected in the SDS-PAGE because the genes encoding for the rest of the proteins are within the same plasmid and open reading frame, further validation will be needed to verify their expression. Fig 3. did not show PhaC (63.4kDa) from pet29b(+)-phaCJ, PhaC (65.1kDa) and PhaB (27.2 kDa) from pet29b(+)-phaCBA. However, because these genes all have their own 6xHis tag, protein purification may be performed to eliminate excess bands to isolate the proteins of interest for a future SDS-PAGE. In addition, a Western Blot can be used to further confirm the protein production.

#### PHB synthesis

The results showed that PHB was produced by cells containing the *phaCBA* operon and the *phaC1J4* biobrick, as evidenced by the presence of white powder after a sodium hypochlorite PHB extraction from cells containing both the *phaCBA* and *phaCJ4* constructs (Fig 6). As discussed below, this white powder was confirmed to be PHB by HPLC analysis. However, the mass of white powder produced by 10 mL of *E. coli* cultures was negligible, and while we could visually confirm PHB production, there was no way to accurately gauge the mass of PHB produced during the lab-scale tests which were conducted. Future tests would need to use more sensitive methods of PHB quantification or work with larger volumes of *E. coli* cultures.

**Fig 6.**
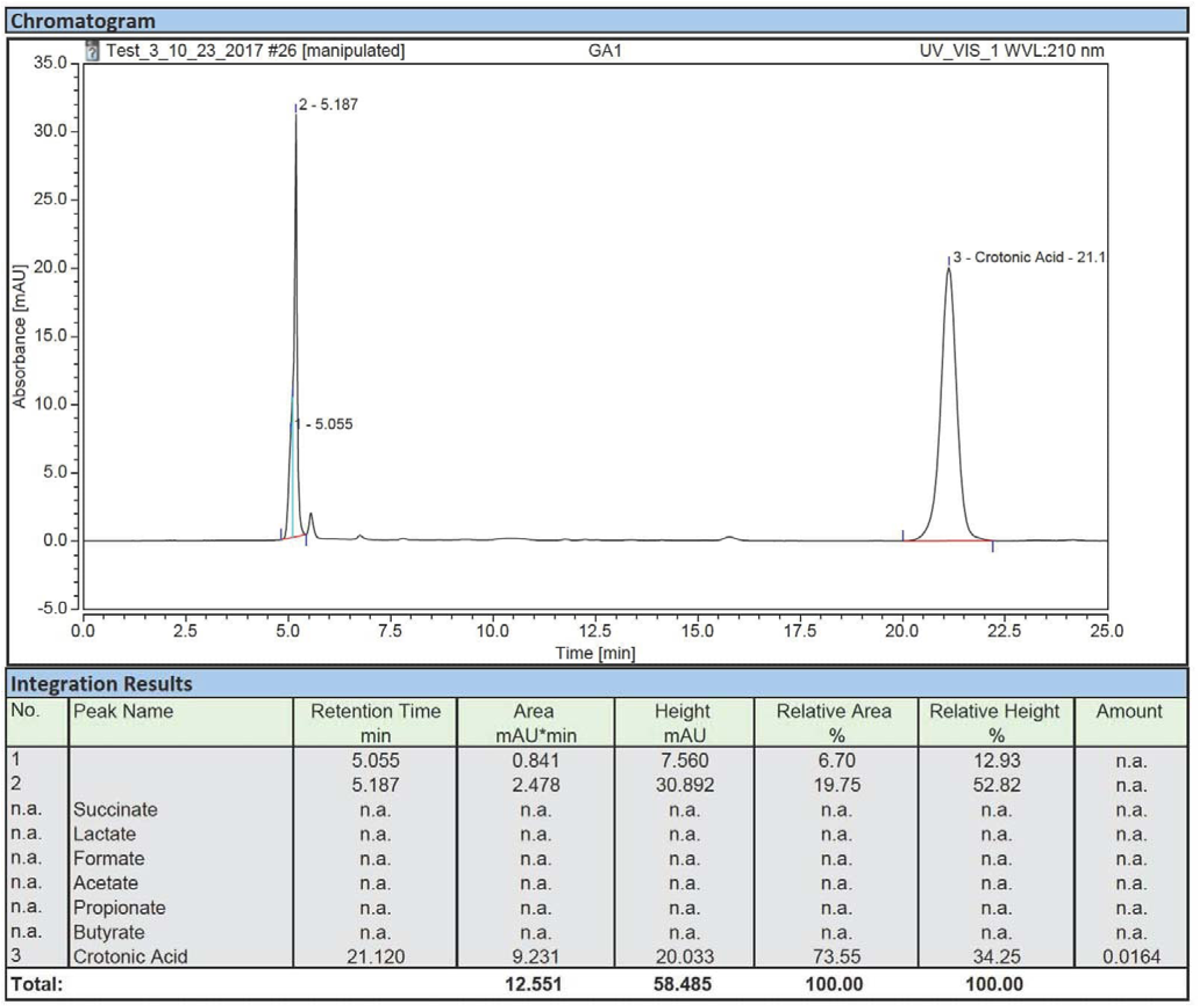
HPLC results of sample containing PHB digested with sulfuric acid. HPLC analysis of sample obtained from sulfuric acid digestion of product extracted from *E. coli* (BL21) transformed with *phaCBA* construct. Product was digested for 30 mins and a dilution factor of 32 was used.

#### HPLC Analysis -*phaCBA* and *phaC1J4*

Samples obtained through chemical extraction of *E. coli* transformed with our genetic constructs were digested with sulfuric acid, and the resulting digest was analyzed using HPLC. HPLC analysis of the sample from *E. coli* transformed with pET29b plasmid containing *phaCBA* operon showed that 0.0282006 mM of crotonic acid was produced when a dilution factor of 32 was used and product digested for 30 mins. HPLC analysis of the digested product obtained from sulfuric acid digestion of sample when *E. coli* transformed with pET29b plasmid containing *phaCJ* operon was used also indicated that crotonic acid was present. Hence, the samples obtained using the two genetic constructs contained PHB. However, the amount of PHB in the sample could not be determined because the amount of crotonic acid detected was too low. The low concentration of crotonic acid could have been due to a number of reasons. Long duration of digestion of sample could have resulted in degradation of crotonic acid. In order to avoid degradation, an optimal digestion time needs to be selected by conducting HPLC with various digestion times. The method of PHB extraction could also be optimized to minimize the loss of PHB. Nonetheless, HPLC was successful in confirming production of PHB, but due to a number of limitations the amount of PHB in the samples could not be determined.

#### Secretion

The results show that 48 hours after induction initiation there was a 114% increase in the amount of PHB secreted by cells that contain *phaCAB-phasin-HlyA* compared to control cells, which contain pSB1C3-phaCAB only. However, there is very little difference between the two groups at 24 hours (Fig 10). Therefore it can be inferred that PHB secretion with phasin-HlyA occurred only after at least 24 hours the induction of cultures with IPTG. These results demonstrate that the type I secretion mechanism in *E. coli* can successfully secrete PHB when coupled with the phasin molecule from *R. eutropha*. The production of "secreted" PHB in control cells may not be PHB that was actually secreted, but instead it may be PHB that was released into the media as a result of cell death and lysis, which is a normal part of the *E. coli* life-cycle. Rahman et. al obtained similar results in their 2013 study on the secretion of PHB in *E. coli*, in which an initiation of PHB secretion between 24 and 48 hours after induction was observed [9].

## Process Development

### Process Overview

The proposed process that converts astronauts’ fecal matter to a bioplastic that can be used to 3D print tool on Mars consists of four major steps (Fig 14). In the first step, astronauts’ feces are collected into a storage tank using a vacuum toilet. Feces are then transferred to a fermenter for 3 days, where natural gut flora produces VFA. In the second step, the liquid containing VFA along with other nutrients found in human feces are separated from solid particles using centrifugation followed by a self-cleaning filter. The obtained liquid is passed to another storage tank. In the third step, the supernatant containing VFA is added to a fermenter, which is then inoculated with PHB-producing and PHB-secreting *E. coli.* In the fourth and last step, PHB secreted by bacteria is extracted from the liquid harvest stream using dissolved air flotation. After drying, the PHB can be used in a Selective Laser Sintering (SLS) 3D printer without additional processing [14]. Each step of the process is further explained in subsequent sections.

**Fig 14.**
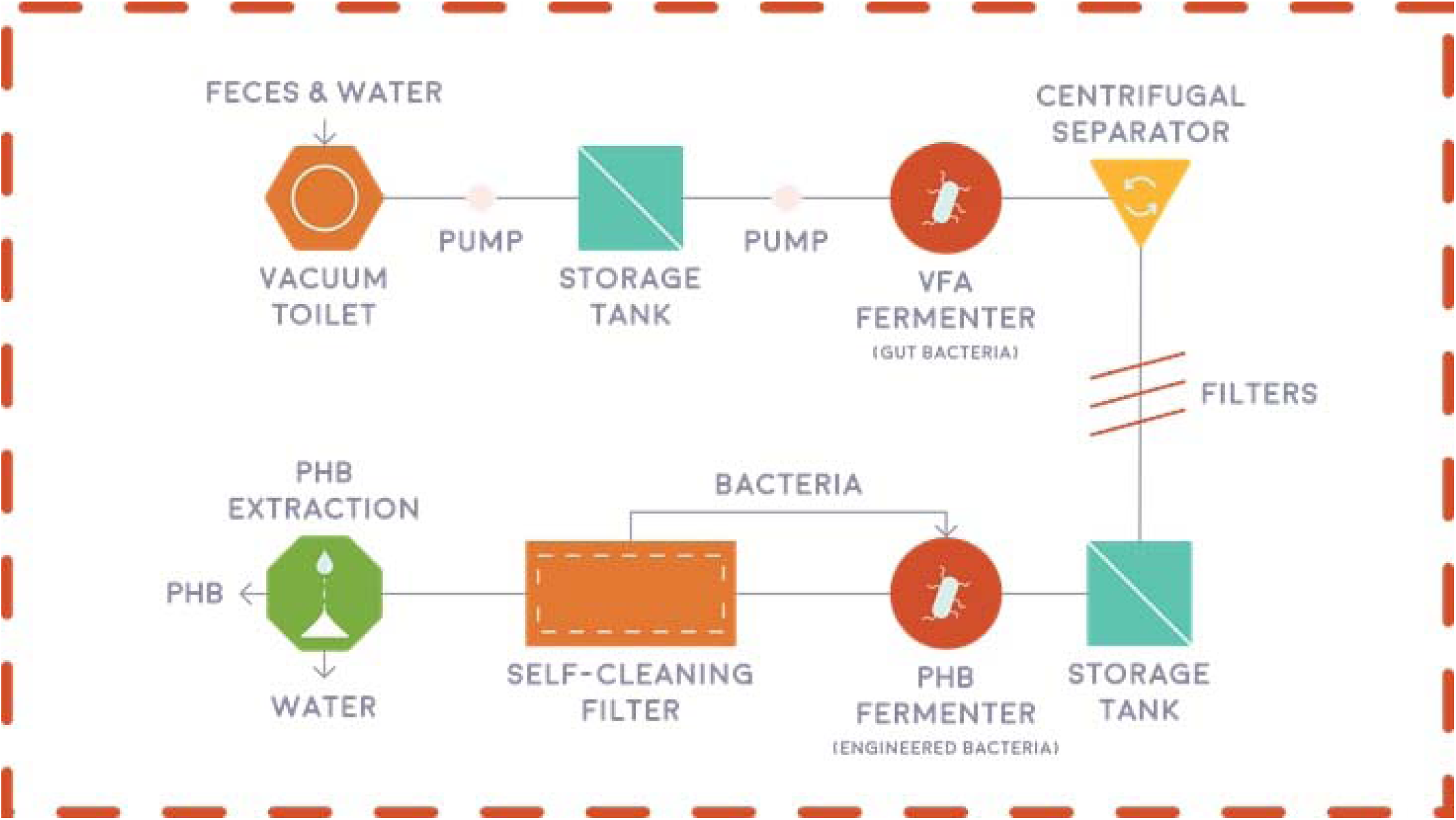
Overview of the proposed process that starts with astronauts’ feces and end with a bioplastic product that can be used to 3D print tool for astronauts.

### VFA Fermentation

In this step, astronauts’ feces are collected into a storage tank using a vacuum toilet. Although the vacuum toilet requires about 0.5 L of water, which is a valuable resource on Mars, per flush, the used water can be recovered at the end of the process using a water processing unit currently used on the International Space Station. The storage tank was designed based on NASA requirements for a fecal collection system, which must be able to collect an average of 150 mL of fecal matter per defecation for 2 defecations per crew member per day [21]. Additionally, the system must be designed to contain 1.5 L of diarrhea discharge. To meet the requirements of our process, the storage tank should also be able to hold 3 days’ worth of fecal matter. Taking these requirements into account with an additional 0.5 L of water entering the storage tank per flush and a crew of 6 astronauts, the volume of the storage tank was found to be 15 L. After the storage tank, feces are fermented in another vessel with naturally occuring bacteria for 3 days at 22°C (room temperature) to increase the concentration of VFA. The fermenter will use an agitator to prevent the settling of solids and bacteria during fermentation. Temperature, pH, and dissolved oxygen will not be controlled.

### VFA Extraction

In this step, the liquid is recovered from feces and separated from solid particles to obtain a sterile, VFA-rich stream for the PHB fermentation step. Large solids are removed using centrifugation and smaller solids along with bacteria are then removed using a self-cleaning filter.

Alternative designs considered for this step included torrefaction and a screw press dewatering system. Torrefaction is defined as thermochemical treatment of biomass in the absence of oxygen and can recover both natural and pyrolytic water. While the torrefaction process was a better alternative based on ESM, the liquid product of torrefaction containing water and VFA was too acidic and lacked required nutrients to support bacterial growth. Centrifugation followed by filtration recovered sufficient amounts of water and had a lower ESM value than a screw press dewatering system. However, we propose torrefaction treatment for the sludge remaining after centrifugation. Torrefaction treatment would recover additional water and produce char, which can be used as a building material, radiation shielding or as a food substrate.

### PHB Fermentation

In this step, VFA-rich stream is fermented with PHB-producing *E. coli* in a continuous process. The process occurs in a 5-L stirred-tank bioreactor at 37°C under anaerobic conditions. Continuous fermentation is achieved using a self-cleaning filter with 0.2 um pores, which separates bacteria from the liquid harvest stream and recycles bacteria back into the bioreactor (Fig 15). The resulting bacteria-free liquid harvest stream contains PHB secreted by bacteria, dissolved salts, and unused volatile fatty acids.

**Fig 15.**
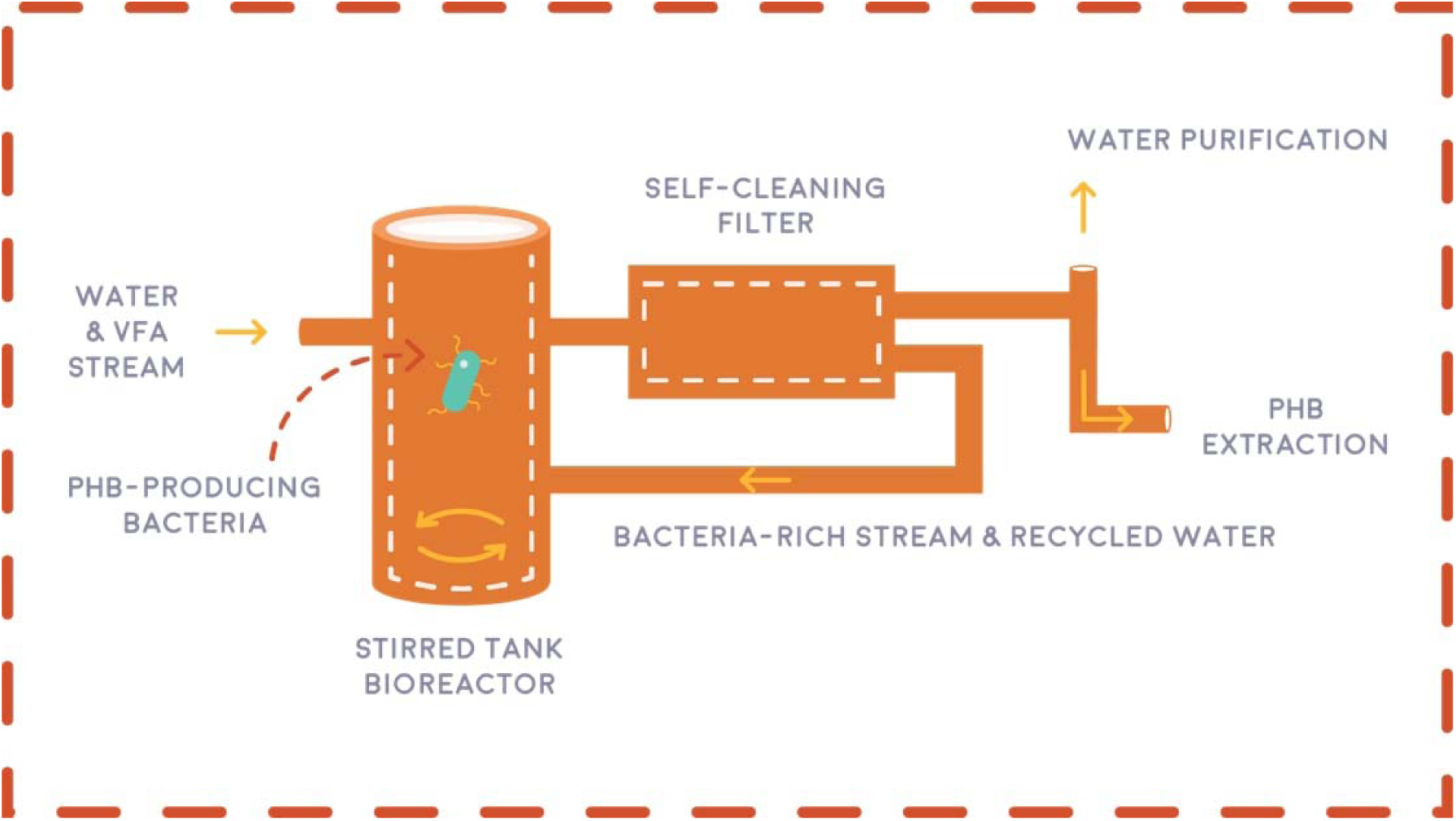
Overview of the proposed PHB fermentation process, where PHB is continuously produced using engineered bacteria.

To achieve a continuous fermentation system, alternative designs considered for this step included various configurations of membrane bioreactors. However, the major disadvantage of membrane bioreactors is fouling, resulting in high crew time requirements and transportation costs due to consumables.

### PHB Extraction

Extraction of PHB nanoparticles from the liquid harvest stream is achieved using dissolved air flotation (DAF) (Fig 16). In DAF, air saturated water is injected into the vessel containing PHB particles suspended in liquid. The resulting air bubbles capture PHB nanoparticles and float up, resulting in accumulation of PHB at the top of the liquid. This layer of PHB can then be removed and dried to obtain the final product.

Alternative designs considered for this step included chemical coagulation and electrocoagulation. While chemical coagulation would require shipment of chemicals to Mars, electrocoagulation resulted in accumulation of sludge along with PHB during laboratory experiments. Dissolved Air Flotation was therefore selected as the preferred design option.

**Fig 16.**
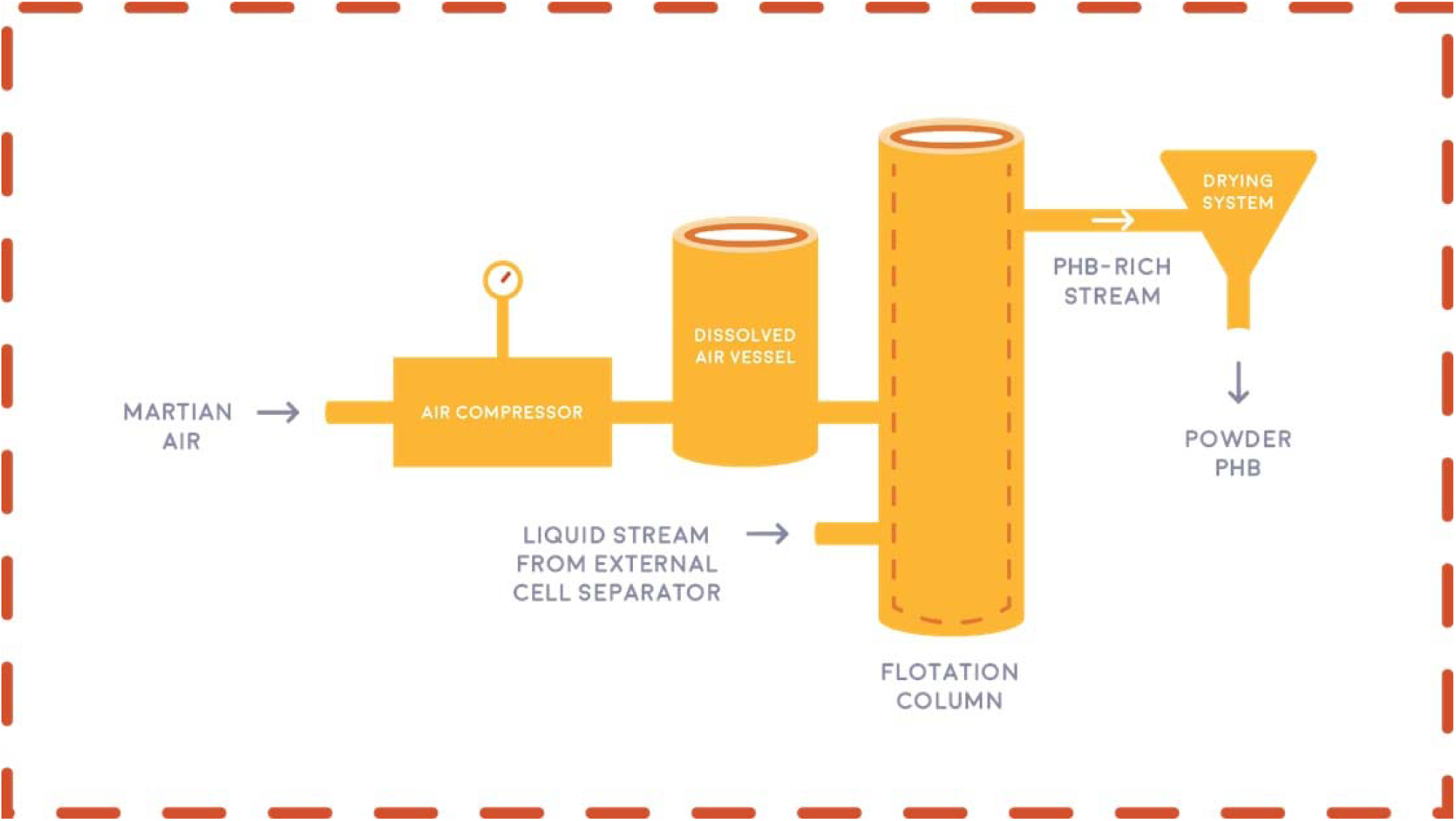
Overview of the proposed PHB extraction method using dissolved air flotation.

### Applied Design

When developing the physical process on Mars that starts with feces and ends with the bioplastic product, we identified design requirements based on advice from experts. Then, we iterated the design of our process based on feedback from space industry professionals including Canadian astronauts Dr. Robert Thirsk and Col. Chris Hadfield, Dr. Matthew Bamsey (Chief Systems Engineer at the German Aerospace Center), and Dr. Pascal Lee (Principal Investigator of the Haughton-Mars Project at NASA Ames Research Center and a co-founder of the Mars Institute). Furthermore, we used ESM analysis, a method recommended by NASA specifically for advanced life support systems, to evaluate the feasibility of our process. Additionally, we investigated applications for solid human waste that NASA is considering and whether the byproducts of our process have any useful applications. These considerations would allow us to integrate our process with NASA’s systems. Lastly, we considered the safety of our design as it was identified as one of the most important criteria based on interviews with experts.

## Conclusion & Future Directions

The key aims of this project were to optimize PHB production in *E. coli*, to optimize PHB secretion from *E. coli* to the surrounding media, and to design a start-to-finish process for PHB production from human waste on a potential future Mars base. Further testing needs to be done to definitively claim that the first two aims were accomplished.

The third aim (the development of the PHB production/ waste management process for Mars bases) has been partially achieved, with all components of the process accounted for and several components confirmed to work at the lab scale.

The dissolved air flotation aspect of the PHB production process will be further developed as a part of the Canadian Reduced Gravity Experiment Challenge (Can-RGX), which will take place in the summer of 2018. As a part of Can-RGX, the dissolved air flotation aspect of our project will be optimized for microgravity in order for PHB production to occur en route to Mars (as well as on the Martian surface).

## Acknowledgements

Dr. Mayi Arcellana-Panlilio – the primary supervisor of iGEM Calgary 2017. Without her guidance and support, the project would not have been realized.

Rachelle Varga – a Teaching Assistant for iGEM Calgary 2017. She assisted with laboratory discussions, as well as supervision, and provided the team with applicable protocols.

David Feehan – a Teaching Assistant for iGEM Calgary 2017. He provided the team with supervision and advice on laboratory techniques.

Dr. Richard Moore -provided the transformation protocol, which which was used to transform the *E. coli* BL21 (DE3) and *E. coli* DH5α used in the experiments.

Dr. Elke Monika Lohmeier-Vogel – gave feedback on the progress of the project and helped research the biochemical pathways of PHB production with the team.

Dr. Craig Jenne – advised the team on the safety aspects of the project on the users and the surrounding environment on Mars.

Dr. Fabiola Aparicio-Ting – guided us on the ethical and societal considerations of the project. Brad Prince – Provided the cost analysis of PHB production using synthetic biology.

Dr. Nashaat Nassar – helped with process development and suggested the use of electrical coagulation for PHB extraction.

Anirban Chakraborty and Jayne Rattray – trained the team on using the HPLC and helped set up runs and methods, as well as troubleshooting, for VFA and PHB quantification.

Author Contributions
Conceptualization: KS XC AK PG MAN HWC SI ZW AI MO JG PL LB BS RCV MAP Methodology: AK KS XC SI MAN HWC JG AI PG ZW LB PL MO BS Software: MO Formal Analysis: AK MAN AI Investigation: KS XC PG SI AK HWC MAN ZW LB JG PL MO RCV DF BS Resources: MAP RCV AK DF Writing Original Draft Preparation: XC SI AK ZW PG KS MAN HWC JG LB Writing – Review & Editing: XC AK SI ZW MAP LB HWC Visualization: TL MO XC AK HWC Supervision: MAP RCV DF Project Administration: MAP RCV DF AK XC SI KS Funding Acquisition: MAP RCV AK KS XC PG SI MO PL ZW LB MAN HWC BS JG

